# *In vivo* exposure of mixed microplastic particles in mice and its impacts on the murine gut microbiome and metabolome

**DOI:** 10.1101/2025.07.15.664901

**Authors:** Kyle Joohyung Kim, Marcus Garcia, Aaron S. Romero, Yan Jin, Jinhua Chi, Matthew J. Campen, Haiwei Gu, Jason R. Richardson, Eliseo F. Castillo, Julia Yue Cui

## Abstract

Microplastics (MPs) are emerging environmental contaminants due to increasing global plastic production and waste. Microplastics, defined as plastic particles less than 5 mm in diameter, are formed through degradation of larger plastics via sunlight, weathering, and microbes. These plastic compounds are widely detected in water, soil, food, as well as human stool and blood. The gut microbiome, often referred to as our second genome, is important in human health and is the primary point of contact for orally ingested microplastics. To investigate the impact of ingested MPs on the gut microbiome and the metabolome, 8 weeks-old male and female C57/BL6 mice were orally gavaged mixed plastic (5 um) exposure consisting of polystyrene, polyethylene, and the biodegradable/biocompatible plastic, poly-(lactic-co-glycolic acid) twice a week for 4 weeks at 0, 2, or 4 mg/week (n = 8/group). Fecal pellets were collected for bacterial DNA extraction and metagenomic shotgun sequencing, and serum was subjected to targeted and untargeted metabolomics. MPs exposure resulted in significant sex-specific and dose-dependent changes to the gut microbiome composition along with substantial regulation of the predicted metabolic pathways. Untargeted metabolomics in serum showed that a low MPs dose displayed a more prominent effect on key metabolic pathways such as amino acid metabolism, mitochondrial function, and inflammation. Additionally, SCFA-targeted metabolomics showed significant changes in neuroprotective SCFAs levels in both sexes by MPs exposure. In conclusion, our study has demonstrated that microplastics dysregulate the gut microbiome and serum metabolome, providing critical insights into potential human disease risks associated with microplastic contamination.

## Introduction

Since the 1950s, global plastic production has surged (UN Environment Programme, 2024), with more than 430 million tons of plastic produced worldwide annually (Galloway et al. 2017). This widespread use of plastic products contributes significantly to the accumulation of plastic waste in the environment, increasing human exposure to plastic byproducts. Plastic debris, subjected to natural factors such as UV radiation, erosion, and microbial degradation, breaks down into smaller microplastics (MPs) (Papini et al. 2024; Zettler et al. 2013) that are defined as particles with a diameter of 5 mm (5000 μm) or less, with those smaller than 1 μm categorized as nanoplastics. Their pervasive presence in various settings—including soil, household dust, and aquatic environments raises significant environmental and public health concerns (Papini et al. 2024). Moreover, microplastics have been detected in food items such as fruits, vegetables, and seafood (Aydin et al. 2023; Jin et al. 2021), which highlights an emerging public health concern regarding the toxicity of microplastics to both humans and wildlife.

Microplastics can enter the human body through ingestion, inhalation, and dermal absorption (Cverenkárová et al. 2021; Campanale et al. 2020; Enyoh et al. 2020). Ingestion is the most common route, occurring through contaminated food and water, as well as non-dietary ingestion from various environmental sources (Sun and Wang 2023). Due to their small size and high abundance, microplastics can readily enter and become bioavailable within the mammalian gastrointestinal (GI) tract and have been found in numerous tissues, including the brain (Nihart et al. 2025). Research suggests that on average, an adult may ingest between 0.1 to 5 grams of microplastic particles per week (Senathirajah et al. 2021). Studies have demonstrated the bioaccumulation of microplastic particles in the 5-20um diameter within the gut tissue of PE microsphere exposed mice (Yang et al. 2019). In Sprague-Dawley rats, the accumulation of polystyrene microspheres measuring 0.5 μm and 1 μm has been observed in the stomach and intestinal cavities, respectively (Reineke et al. 2013).

Following accumulation in the GI tract, microplastics may disrupt gut epithelium and compromise gut barrier integrity, including tight junctions, potentially increasing the risk of systemic exposure to harmful substances and microbial metabolites (Lee et al. 2018). Studies in rats exposed to polystyrene microspheres have shown decreased transcripts of tight junction proteins within the gut epithelium, indicating potential barrier disruption (Huang et al. 2023). Additionally, murine studies exposed to polystyrene microspheres have demonstrated physical damage within the intestinal wall after prolonged exposure (Xiao et al. 2022).Beyond compromising gut barrier integrity, microplastics have been implicated in causing inflammation and ulceration of the GI tract (Yan et al. 2022; Zolotova et al. 2023). These effects can exacerbate existing conditions such as inflammatory bowel diseases (IBD), including Crohn’s disease and ulcerative colitis, by increasing intestinal permeability, inflammation, and susceptibility to infections (Yan et al. 2022; Zolotova et al. 2023).

The gut microbiota, which plays a crucial role in maintaining gut epithelial integrity and immune homeostasis, can also be altered by microplastic exposure (Ndoul 2020). Dysbiosis of gut microbiota has been associated with various conditions including inflammatory bowel diseases, colon cancer, and neurological disorders mediated by the gut-brain axis (Menees 2018; Kim 2021; Tiwari et al. 2023). Mice exposed to polystyrene microspheres demonstrated significant alterations in gut microbiome composition and associated microbial transcriptome, highlighting the potential for microplastics to influence gut health indirectly through microbiota modulation (Lu et al. 2018).

While the influence of microplastics on gut health and the gut microbiome is increasingly recognized, many questions remain regarding the specific mechanisms of toxicity within the gut and their indirect effects on other organ systems such as the brain. Addressing these knowledge gaps is crucial for understanding the full scope of health risks associated with microplastic exposure and developing effective mitigation strategies.

Therefore, this study aims to investigate compositional changes in the murine gut microbiome and associated metabolic signatures in blood serum following exposure to microplastic particles. By elucidating these effects, the study seeks to contribute to a deeper understanding of the impacts of microplastics on gut health and their potential implications for systemic health, including neurological function.

## Methods

### Animal Husbandry and Dosing

Male and female C57BL/6 wild-type mice (8-12 weeks of age at the beginning of the study) were obtained from Taconic Biosciences (Rensselaer, New York). Animals were housed in an Association for Assessment and Accreditation of Laboratory Animal Care (AAALAC)-approved facility at the University of New Mexico Health Sciences Center. Animals were maintained at constant temperature (20–24°C), relative humidity (30%–60%), and on a 12-h light/dark cycle throughout the study. Animals had *ad libitum* access to standard rodent chow and water. All experiments were approved by the Institutional Animal Care and Use Committee of the University of New Mexico Health Sciences Center (Protocol no. 23-201398-HSC) and by the National Institutes of Health guidelines for using live animals. The American Association for Accreditation of Laboratory Animal Care accredits the University of New Mexico Health Sciences Center. Mice were exposed to a mixture of microplastics (MPs) via oral gastric gavage over four weeks at 0 mg/week (0.1% tween 20, male n=4, female n=4), 100ul of 10mg/ml per dose twice a week equating to 2 mg/week (male n=4, female n=4), and 200ul of 10mg/ml per dose twice a week equating to 4 mg/week (male n=4, female n=4) of 5 µm microspheres from Degradex® Phosphorex (Garcia et al. 2024). The MPs mixture consists of polystyrene (PS), polyethylene (PE), and poly (lactic-co-glycolic acid) (PLGA) microspheres at a 1:1:1 ratio. Before oral gavage, 5 µm polystyrene, polyethylene, and poly (lactic-co-glycolic acid) microspheres were mixed to a 1:1:1 ratio concentration and stored per the manufacturer’s recommendations in a 10 mg/ml concentration at 4°C.

### Metagenomic shotgun sequencing and data analysis

Fecal pellets were collected from C57/BL6 mice after 4 weeks of MPs exposure. Microbial DNA was extracted using an O.M.E.G.A EZNA Stool DNA extraction kit (OMEGA Biotech Inc, Norcross, Georgia). DNA concentration was quantified using a Qubit fluorometer (Thermo Fischer Scientific, Waltham, Massachusetts).

Shallow metagenomic shotgun sequencing was performed on the Illumina NovaSeq 6000 sequencer utilizing the BoosterShot sequencing method (Diversigen, New Brighton, Minnesota). The sequencing was performed with 2×150 bp paired end reads, achieving a minimum of 2 million reads per sample. DNA sequences were aligned to a curated database containing all representative bacterial genomes, including additional manually curated mouse-specific Metagenomically Assembled Genomes (MAGs) and cell-cultured genomes. Only high-quality MAGs (completeness > 90% & contamination < 5% via checkM) were included to ensure quality. Alignments were performed with a 97% match-identity against all reference genomes, with each input sequence compared to every reference sequence in the Diversigen DivDB-Mouse database. Ties were resolved by minimizing the overall number of unique operational taxonomic units (OTUs). Taxonomy assignment was based on the lowest common ancestor consistent across at least 80% of all reference sequences tied for the best hit according to the Genome Taxonomy Database (GTDB r95). (Diversigen, New Brighton, Minnesota).

Samples with fewer than 10,000 detected sequences were excluded from downstream analysis. OTUs representing less than one-millionth of all strain-level markers and those with less than 0.01% of their unique genome regions covered (and < 0.1% of the whole genome) at the species level were also discarded. The number of counts for each OTU was normalized to the genome length of the OTUs. Strain coverage maps were established using filtered OTU, stratified from the overall OTU general table.

Bray-Curtis beta diversity indices were calculated using the filtered taxonomy and KEGG modules/enzymes tables through QIIME 1.9.1. (Navas-Molina et al. 2013). Shannon index alpha diversity indices were established using rarefied and filtered taxonomy table sets (with a minimum sequence depth of 10,000 reads) and were used through QIIME 1.9.1

Normalized count data was processed using Rstudio, and differentially regulated microbes, pathways, and modules were identified using EdgeR (Robinson et al. 2010). The trimmed mean of M (TMM) normalization and Bonferroni’s Correction were applied, with additional filtering criteria described as follows:

1. Taxa: fold change > 1.3, *p*-value < 0.05
2. Modules: fold change > 2.0, *p*-value < 0.05
3. Enzymes: fold change > 2.0, *p*-value < 0.01
4. L3 (class level) pathways: fold change > 2.0, p-value < 0.01).

Significance lists were further filtered using various row-means average counts significance cut-offs for each data set taxa > 10, modules > 100, enzymes > 400, and L3 pathways > 100). After processing the normalized counts data through edgeR, a file containing the significant data hits is generated for visualization. For data plotting and visualization, ggplot2 (v.3.3.6) and **SigmaPlot (Graffiti LLC. 2022) were utilized for bar plots while heatmaps were plotted using the** ComplexHeatmaps (v.2.13.1) Rstudio package.

### Untargeted LC-MS Metabolomics

Acetonitrile (ACN), methanol (MeOH), ammonium acetate, and acetic acid, all LC-MS grade, were purchased from Fisher Scientific (Pittsburgh, PA). Ammonium hydroxide was bought from Sigma-Aldrich (Saint Louis, MO). DI water was provided in-house by a Water Purification System from EMD Millipore (Billerica, MA). PBS was bought from GE Healthcare Life Sciences (Logan, UT). The standard compounds corresponding to the measured metabolites were purchased from Sigma-Aldrich (Saint Louis, MO) and Fisher Scientific (Pittsburgh, PA).

Frozen serum samples were first thawed overnight under 4°C, and 50 μL of each sample was placed in a 2 mL Eppendorf vial. The initial step for protein precipitation and metabolite extraction was performed by adding 500 μL MeOH and 50 μL internal standard solution (containing 1,810.5 μM ^13^C_3_-lactate and 142 μM ^13^C_5_-glutamic acid). The mixture was then vortexed for 10 s and stored at -20°C for 30 min, followed by centrifugation at 14,000 RPM for 10 min at 4° C. The supernatants (450 μL) were collected into a new Eppendorf vial and dried using a CentriVap Concentrator (Labconco, Fort Scott, KS). The dried samples were reconstituted in 150 μL of 40% PBS/60% ACN. A pooled sample, which was a mixture of all plasma samples was used as the quality-control (QC) sample.

The untargeted LC-MS metabolomics method used here was modeled after that developed and used in a growing number of studies (Gu et al. 2015; Wei et al. 2021; Jin et al. 2023; Mohr et al. 2024; Scieszka et al. 2023; Kim et al. 2023; Garcia et al. 2024). Briefly, all LC-MS experiments were performed on a Thermo Vanquish UPLC-Exploris 240 Orbitrap MS instrument (Waltham, MA). Each sample was injected twice, 10 µL for analysis using negative ionization mode and 4 µL for analysis using positive ionization mode. Both chromatographic separations were performed in hydrophilic interaction chromatography (HILIC) mode on a Waters XBridge BEH Amide column (150 x 2.1 mm, 2.5 µm particle size, Waters Corporation, Milford, MA). The flow rate was 0.3 mL/min, the auto-sampler temperature was kept at 4 °C, and the column compartment was set at 40 °C. The mobile phase was composed of Solvents A (10 mM ammonium acetate, 10 mM ammonium hydroxide in 95% H_2_O/5% ACN) and B (10 mM ammonium acetate, 10 mM ammonium hydroxide in 95% ACN/5% H_2_O). After the initial 1 min isocratic elution of 90% B, the percentage of Solvent B decreased to 40% at t=11 min. The composition of Solvent B maintained at 40% for 4 min (t=15 min), and then the percentage of B gradually went back to 90%, to prepare for the next injection. Using a mass spectrometer equipped with an electrospray ionization (ESI) source, we will collect untargeted data from 70 to 1050 m/z.

To identify peaks from the MS spectra, we made extensive use of the in-house chemical standards (∼600 aqueous metabolites), and in addition, we searched the resulting MS spectra against the HMDB library, Lipidmap database, METLIN database, as well as commercial databases including mzCloud, Metabolika, and ChemSpider. The absolute intensity threshold for the MS data extraction was 1,000, and the mass accuracy limit was set to 5 ppm. Identifications and annotations used available data for retention time (RT), exact mass (MS), MS/MS fragmentation pattern, and isotopic pattern. We used the Thermo Compound Discoverer 3.3 software for aqueous metabolomics data processing. The untargeted data were processed by the software for peak picking, alignment, and normalization. To improve rigor, only the signals/peaks with CV < 20% across quality control (QC) pools, and the signals showing up in >80% of all the samples were included for further analysis.

Raw untargeted metabolomics data were filtered using a CV(QC) value of <20% and filtered data values were normalized to the averaged QC values and one-way analysis of variance (ANOVA) was utilized using Tukey’s posthoc test with a p-value of 0.05 in Rstudio. Pathway analysis data was visualized using the MetaboAnalyst software v.5.0 (https://www.metaboanalyst.ca/MetaboAnalyst/home.xhtml). FDR was set at or below 0.1 and a p-value of 0.05 to identify significant metabolic pathways.

### GC-MS Measurement of SCFAs

Acetic acid was bought from Thermo Fisher Scientific (Fair Lawn, NJ). Propionic acid, isobutyric acid, butyric acid, 2-methylbutyric acid, isovaleric acid, valeric acid, 2-methylpentanoic acid, 3-methylpentanoic acid, isocaproic acid, caproic acid, 2-methylhexanoic acid, 4-methylhexanoic acid, heptanoic acid, hexanoic acid-6,6,6-d_3_ internal standard, N-tert-Butyldimethylsilyl-N-methyltrifluoroacetamide (MTBSTFA), and methoxyamine hydrochloride were purchased from Sigma-Aldrich (St. Louis, MO).

For serum/plasma samples, 20 μL of each sample was mixed with 30 μL aqueous NaOH (0.1M in water), 20 μL IS (hexanoic acid-6,6,6-d3; 200 µM) and 430 μL MeOH in a 1.5 mL Eppendorf tube. The pH value for the mixture was 9. Samples were then vortexed for 10 s and stored at -20 °C for 20 min. After centrifugation at 14,000 RPM for 10 min at 4 °C, 450 μL supernatant was removed into a new Eppendorf tube. The samples were dried under vacuum at 37 °C for 120 min using a CentriVap Concentrator (Labconco, Fort Scott, KS).

Each sample was first derivatized with 40 µL of methoxyamine hydrochloride solution in pyridine (MeOX, 20 mg/mL) under 60 ℃ for 90 min. Next, 60 µL of MTBSTFA was added, and the mixture was incubated under 60 ℃ for 30 min. Then the sample was vortexed for 30 s, followed by centrifugation at 14,000 rpm for 10 min. Finally, 70 µL supernatant was collected into a new glass vial for GC-MS analysis.

GC-MS experiments were performed using an Agilent 7820A gas chromatography system coupled to an Agilent 5977B mass spectrometer (Agilent Technologies, Santa Clara, CA). Chemical derivatives in the samples were separated using an HP-5 ms capillary column coated with 5% phenyl-95% methylpolysiloxane (30 m×250 µm i.d., 0.25 µm film thickness, Agilent Technologies). 1 µL of each sample was injected, and the solvent delay time was set to 5 min. The initial oven temperature was held at 60 ℃ for 1 min, ramped up to 325 ℃ at a rate of 10 ℃/min, and finally held at 325 ℃ for 10 min. Helium was used as the carrier gas at a constant flow rate of 20 mL/min through the column. The temperatures of the front inlet, transfer line, and electron impact (EI) ion source were set at 250 ℃, 290 ℃, and 230 ℃, respectively. The electron energy was -70 eV, and the mass spectral data were collected in the full scan mode (m/z 30-600).

Agilent MassHunter Workstation Software Quantitative Analysis (B.09.00) was used to process the GC-MS data for compound identification, peak picking, and quantification. The signal-to-noise ratio (S/N) was set to S/N=3. The retention time and quantification mass for each SCFA were determined using its chemical standard. The concentrations of SCFAs in biological samples were calculated using the calibration curves constructed from the corresponding SCFA standards.

Short-chain fatty acid data was normalized to the average QC and statistics were done through one-way analysis of variance (ANOVA) using Duncan’s posthoc test with a p-value of 0.05. Boxplot data visualization was done in Rstudio using the ggplot2 (v.3.3.6).

## Results

### Microbiome profiling

As shown in Figure 2A, MPs did not significantly impact the fecal microbiome’s alpha diversity (Shannon average index) in either sex. Beta diversity (Bray-Curtis index) principal component analysis also showed minimal effect of MPs, except for a distinct separation between control and high MPs groups in male mice (Figure 2B). However, the relative number of differentially regulated gut microbes followed a dose-dependent pattern. A higher number of differentially regulated gut microbes is seen in both male-high and female-high groups (Figure 3A-B).

**Figure 1.**
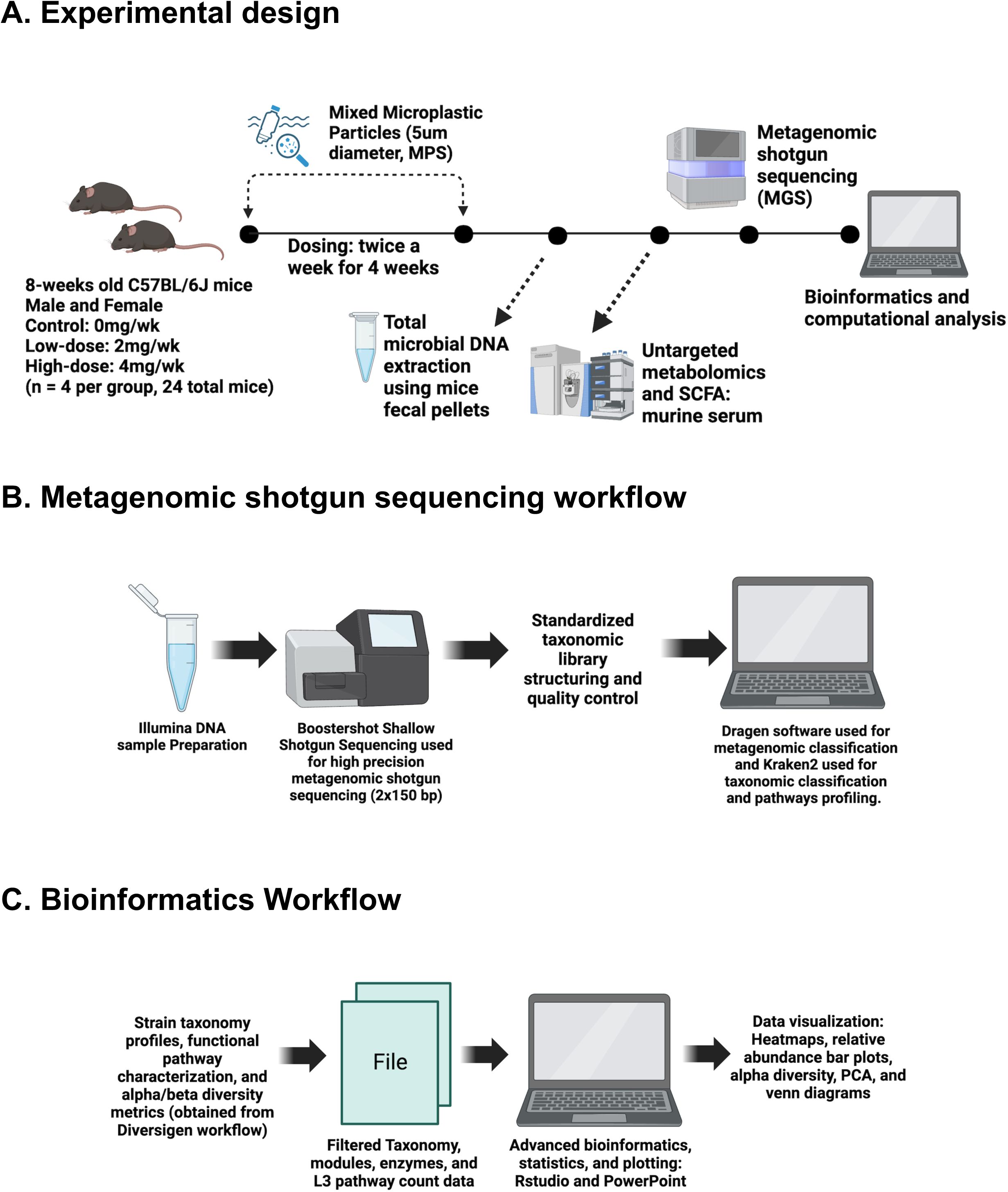
**A.** A general overview of the experimental diagram, 8-week-old male and female C57BL/6 mice were stratified into three exposure groups (control dose of 200ul of 0mg/ml vehicle twice a week, low dose: 100ul of 10mg/ml per dose twice a week for a total of 2mg/week and high dose: 200ul of 10mg/ml per dose twice a week for a total of 4mg/week). At the end of the exposure, fresh fecal pellets were collected for total microbial DNA extraction and subsequent metagenomic shotgun sequencing. Blood serum of the mice was also utilized to conduct both targeted-SCFA and untargeted metabolomics quantification and data were analyzed using advanced bioinformatics and computational analysis. **B.** In the general workflow of metagenomic shotgun sequencing, DNA is inserted into an amplicon sequencer that would conduct a 2×150 bp double-strand DNA sequencing. The raw reads are then matched to a standardized taxonomic library and are structured to fit quality standards for bioinformatic analysis of taxonomic classifications and pathway profiling. **C.** The strain taxonomy profile, functional pathway characterization, and alpha/beta diversity metrics files provided by the sequencing company will then be read through/analyzed using RStudio. Advanced bioinformatics, statistics, and plotting will be conducted using RStudio and PowerPoint.

**Figure 2.**
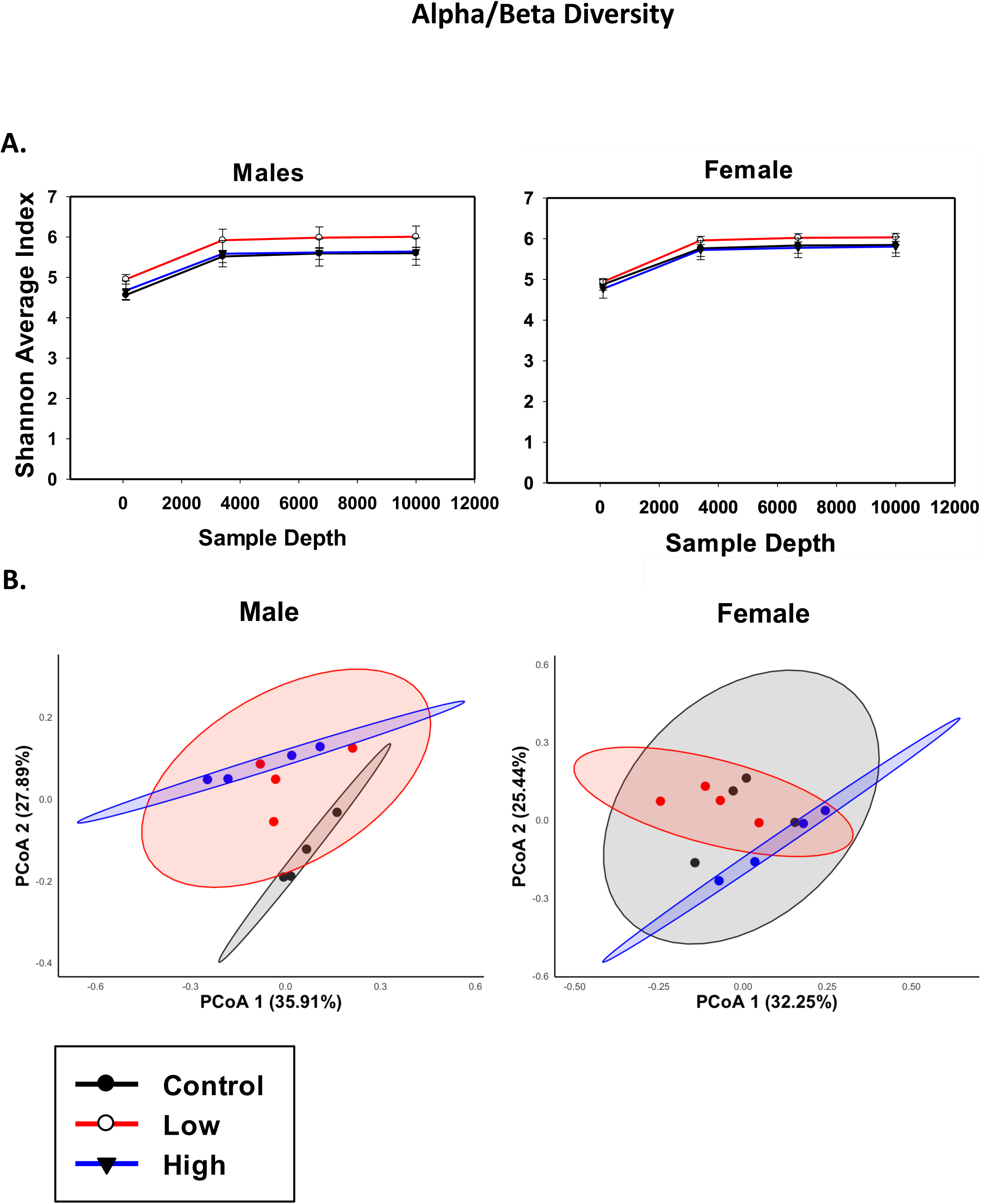
**A.** Alpha diversity (Shannon index) plots reveal that the overall microbial richness in both male and female mice exhibited minimal changes following exposure to mixed microplastic particles (MPs). **B.** beta diversity (Bray-Curtis index) principal coordinate analysis plots showed that in male mice, there was a distinct section between control and high MPs dose exposed groups, whereas such distinction was not prominent in female mice.

**Figure 3.**
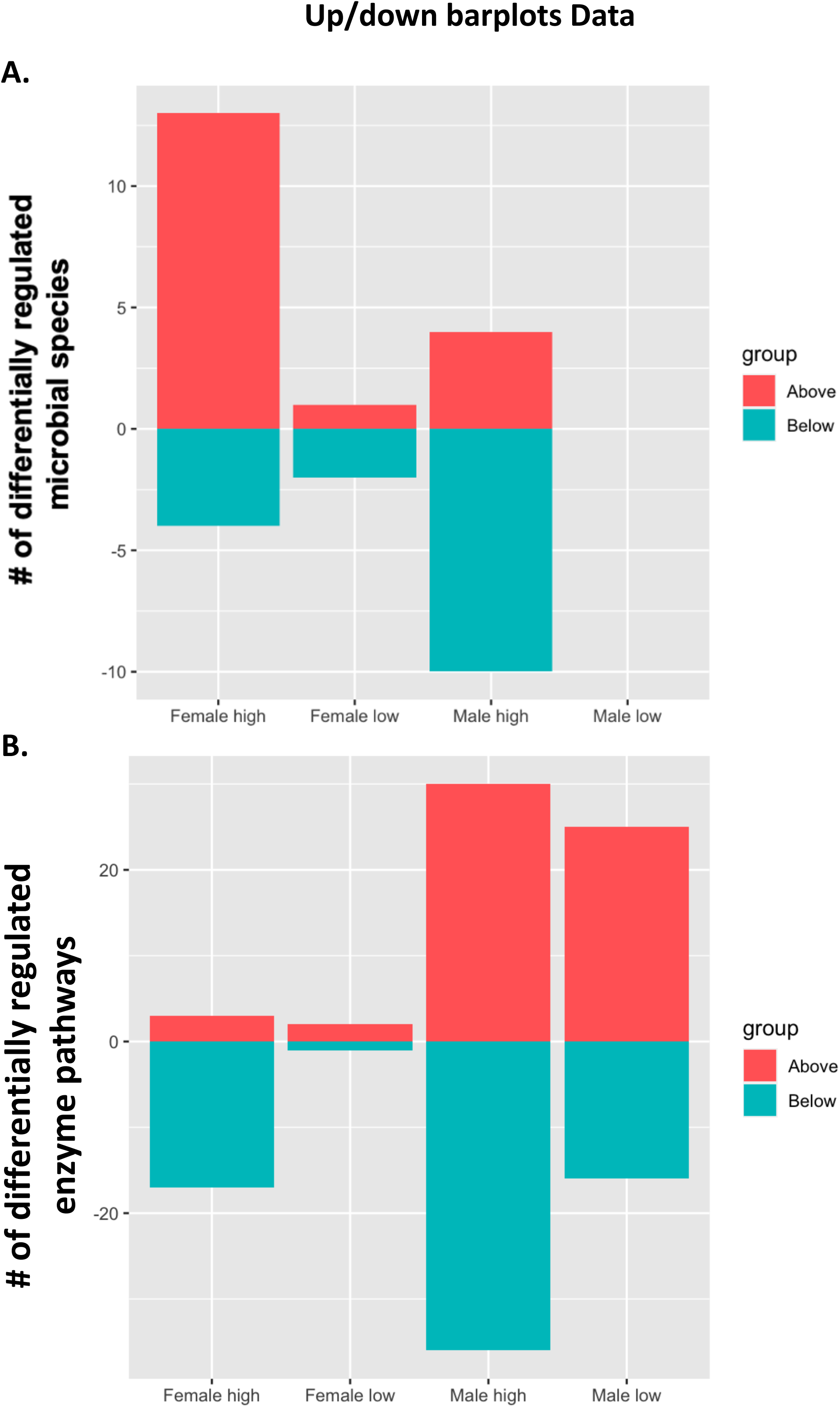
**A.** Bi-directional bar plot showing the number of differentially regulated microbial species, the number of differentially regulated bacterial species was higher in high-microplastic exposed mice groups. The male-high group saw a greater decrease pattern while the female-high group saw a greater increase pattern. **B.** Bi-directional bar plot showing the number of differentially regulated functional enzyme pathways, the male mice group saw a greater number of differentially regulated enzyme pathways compared to female mice group.

Metagenomic shotgun sequencing was conducted in both the vehicle and MPS-exposed male and female mice to determine the effect of MPS exposure on the gut microbiome, as described in MATERIALS and METHODS. The relative abundance (z-scores) of the differentially regulated microbial species are shown in Figure 4A. In male mice, the low MPS dose did not alter any microbes in the gut microbiome, whereas the high MPS dose differentially regulated 14 microbial species, which were partitioned into 2 distinct clusters. The up-regulated cluster by MPS includes *Dubosiella newyorkensis, Dubosiella sp 000403415, COE1 sp002490665*, and *Enterococcus faecalis*; whereas the down-regulated cluster by MPS include *Schaedlerella sp000364245, Rikenella microfusus, Duncaniella sp002492665, Romboutsia timonensis, Romboutsia ilealis, Akkermansia muciniphila, Akkermansia muciniphila A, Akkermansia muciniphila B, Bacteroides xylanisolvens, and Bacteroides thetaiotaomicron*.

**Figure 4.**
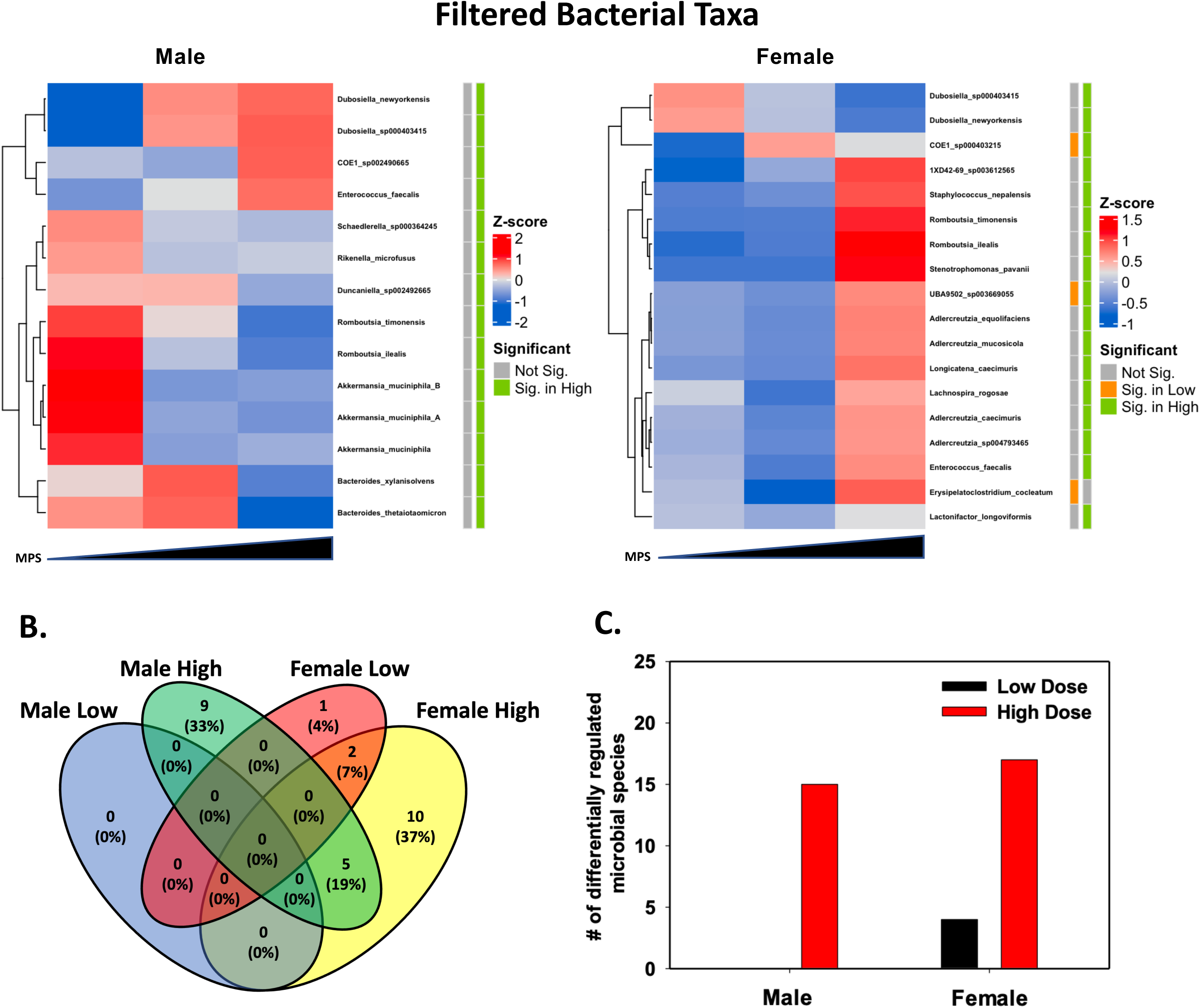
**A.** Heatmaps showing the differentially regulated bacterial species. The number of differentially regulated species of bacteria followed a dose-dependent pattern in both male and female mice male mice saw more differentially decreased bacterial species and female mice had increased bacterial species. **B.** The Venn diagram shows the stratification of the total number of differentially regulated microbial species by sex and treatment, 48% of the differentially regulated bacterial species were seen in female mice alone while 33% of the differentially regulated bacterial species were seen in male mice alone. An overlap of 5 differentially regulated microbes were seen in both male-high and female-high groups which accounted for 19% of the total differentially regulated bacterial species. **C.** A simple bar plot showing the number of differentially regulated bacterial species stratified by sex and treatment. Both male-high and female-high groups saw a similar number of differentially regulated bacterial species but overall, more differentially regulated bacterial species were detected in female mice.

In female mice, the low MPS dose differentially regulated 3 microbes, namely *COE1 sp000403215* (up-regulated), as well as *UBA9502 sp003669055* and *Erysipelactoclostridium cocleatum* (down-regulated). The high MPS dose differentially regulated 18 microbes, which were partitioned into 2 clusters. The up-regulated cluster include *1XD42-69 sp003612565, Staphylococcus nepalensis, Romboutsia timonensis, Romboutsia ilealis, Stenotrophomonas pavanii, UBA9502 sp003669055, Adlercreutzia equolifaciens, Adlercreutzia mucosicola, Adlercreutzia caecimuris, Adlercreutzia sp004793465, Longicatena caecimuris, Lachnospira rogosae, Enterococcus faecalis, Erysipelatoclostridium cocleatum, and Lachonifactor longoviformis.* The down-regulated cluster includes *Dubosiella sp000403415, Dubosiella newyorkensis, and COE1 sp000403215*. To note, while *MPS upregulated COE1 sp000403215* at both doses, the decreased pattern of *UBA9502 sp003669055* and *Erysipelactoclostridium cocleatum* by the low MPS dose was reversed by the high MPS dose.

To further determine the common and unique MPS targeted microbes in different sex and MPS dose groups, a Venn diagram was plotted as shown in Figure 4B. None of the microbes were commonly regulated by MPS in all 4 of the treatment groups. MPS commonly differentially regulated five bacteria in male-high and female-high treatment groups, namely *Dubosiella newyorkensis, Dubosiella sp000403415, Romboutsia ilealis, Romboutsia timonensis,* and *Enterococcus faecalis*. *Dubosiella newyorkensis* and *Dubosiella sp000403415* were upregulated in male-high mice and downregulated in the female treatment groups. *Romboutsia ilealis* and *Romboutsia timonensis* were both downregulated in male-high mice but upregulated in female mice. Only one bacterium, *enterococcus faecalis,* was upregulated in both male-high and female treatment groups.

In both males and females, a dose-response relationship between MPS and the differentially regulated microbes was observed, in that the low MPS dose had no effect in males and minimal effect in females regarding microbes. In contrast, the high MPS dose altered significantly more microbes in both sexes (Fig. 4C).

### Predictive functional analysis (enzymes)

The abundance of the predictive functional microbial enzymes was calculated from the metagenomic shotgun sequencing analysis based on methods established by Diversigen (New Brighton, MN). MPs exposure in the male group resulted in 76 predicted differentially regulated enzymes: 32 predicted enzymes were significantly upregulated, the majority of which were phosphodiesterases, glucosidases, transferases, kinases, primases, synthases, phosphatases, lyases, epimerases, deacetylases, binding cassettes, dehydrogenases, and hydrolases associated with specific amino acid metabolism, bacterial structure, and glycolysis (Fig. S1). 44 predicted enzymes were significantly downregulated, the majority of which were galactosidases, chorismatases, synthases, kinases, transferases, dehydrogenases, catalases, reductases, phosphatases, hydratases, oxidoreductases, and decarboxylases associated with amino acid metabolism, bacterial survivability, and signaling molecule synthesis (Fig. S1).

8 predicted enzymes were significantly regulated in the female high group, namely deacetylases, synthases, glucosidases, and hydrolases associated with amino acid metabolism and sugar metabolism (Fig S1).

A Venn diagram was utilized to visualize the enzyme data set by sex and treatment. There were 30 uniquely predicted regulated enzymes within the malehigh group, namely sulfatases, hydrolases, kinases, dehydrogenases, dehydratases, transferases, phosphatases, reductases, synthases, phosphodiesterase’s, chorismatases, and biosidases associated with glycolysis, bacterial survivability, and bacterial signaling molecule synthesis (Fig 5A.)

**Figure 5.**
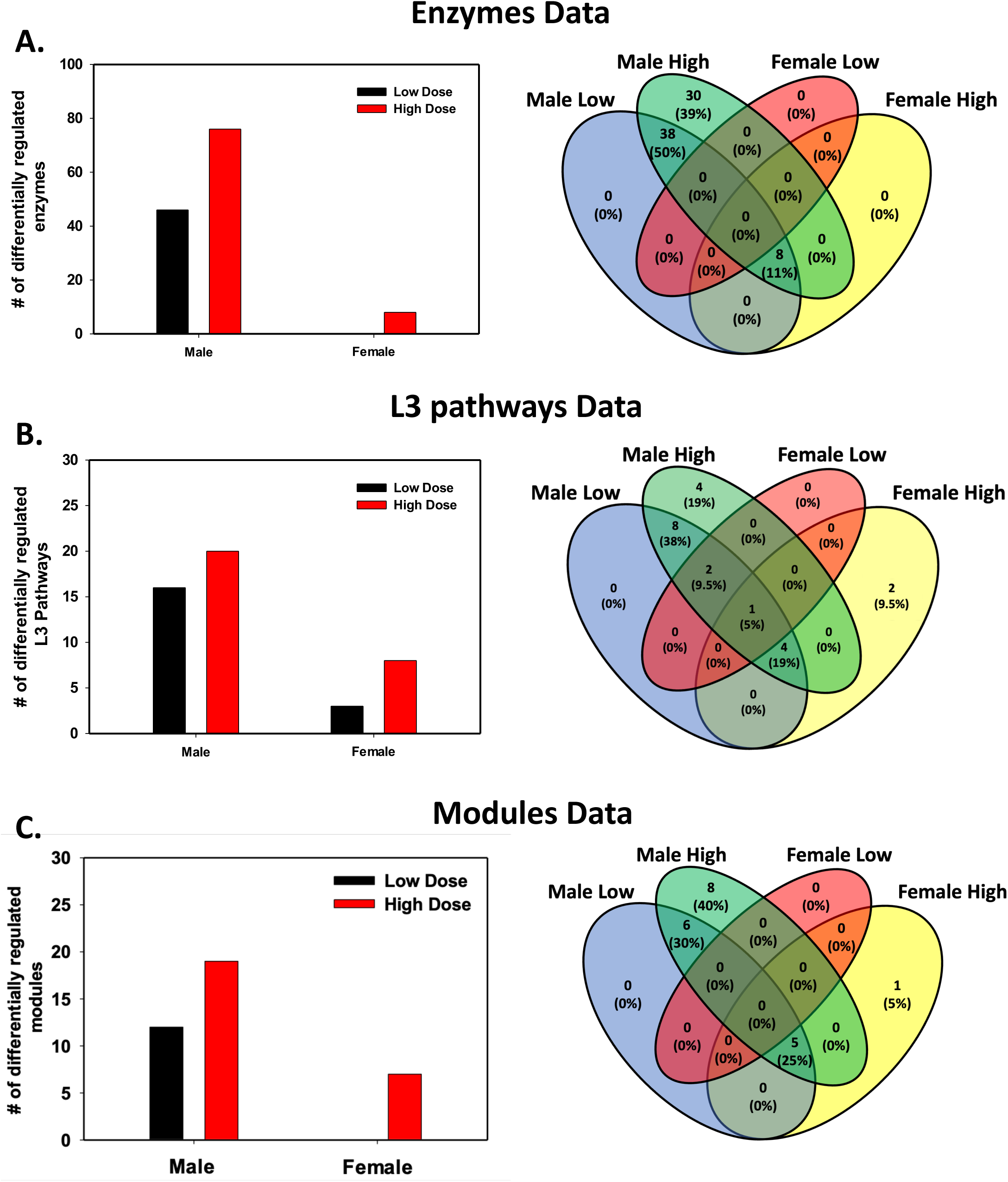
**A.** Venn diagram showing the total number of differentially regulated functional enzymes stratified by sex and treatment, 89% of the differentially regulated functional enzymes were seen only in male mice while 11% of the differentially regulated functional enzymes were shared between the male-high and female-high group. Simple bar plot showing the number of significantly regulated functional enzyme pathways stratified by both sex and treatment. Male groups saw a greater number of differentially regulated functional enzymes compared to the female group. **B.** Venn diagram showing the total differentially regulated L3 functional enzyme pathways stratified by sex and treatment, 54% of the total differentially regulated L3 functional enzyme pathways were associated with only the male group while only 9% of the detected differentially regulated L3 functional enzyme pathways were associated with only females. 37% of the detected pathways were shared between male and female groups. Simple bar plot showing the number of detected differentially regulated L3 functional enzyme pathways stratified by both sex and treatment. A dose dependent trend exists between both male and female groups. Male groups had more significant pathways detected compared to the female group. **C.** 70% of detected modules were specific only to the male group while only 5% of detected modules were specific only to females. 25% of the detected significant modules were shared between both the male and female group. Simple bar plot showing the number of differentially regulated enzyme modules stratified by both sex and treatment. The Male group had a higher number of significantly regulated enzyme modules compared to the female group.

38 predicted enzymes were significantly regulated by MPS exposure in both the male-low and the male-high group, namely: glucosidases, lyases, phosphatases, binding cassettes, galactosidases, binding proteins, oxidoreductases, hydratases, dehydrogenases, peroxidases, synthases, deacetylases, transferases, isomerases, ligases, catalases, hydrolases, decarboxylases, kinases, and primases associated with bacterial cell wall structure, oxidative stress response, and amino acid metabolism (Fig 5 A).

8 predicted enzymes were significantly regulated by MPS exposure in three groups: male-low, male-high, and female-high: glucosidases, synthases, hydrolases, deacetylases, and lysostaphin associated with amino acid metabolism and sugar metabolism (Fig. 5 A).

In both males and females, a dose-response relationship was observed in that low MPs exposure had a weaker effect on the functional predictive enzyme hits compared to high MPs exposure. Male mice had more significantly regulated predicted enzymes associated with MPs exposure, suggesting that the male group was more sensitive toward MPS exposure compared to the female group (Fig. 5 A).

### Predictive functional analysis (L3 pathways)

Significantly regulated L3 (i.e., genus) predictive pathways were analyzed from the metagenomic shotgun sequencing based on methods established by Diversigen (New Brighton, MN). In total, 22 significantly regulated L3 predictive pathways were found among the four treatment groups. 19 L3 predictive pathways were significantly regulated in the male group: 10 L3 predictive pathways were downregulated and associated with peptidases, structural polysaccharides, oxygenase, histidine protein kinases, transporters, cellular structure, polyketide proteins, facilitator proteins, and cell membrane proteins (Fig. S2). 9 predicted pathways were upregulated during MPs exposure in males that were associated with peptidases, cellular structure, oxidoreductases, and cell membrane proteins (Fig. S2). Four predicted pathways were downregulated during MPs exposure in females associated with cell membrane proteins, transporters, and peptidases while five pathways were upregulated in the female group associated with transporters, facilitator proteins, and kinases (Fig. S2).

A Venn diagram was utilized to visualize the unique and shared differentially regulated L3 predictive pathways among the treatment groups. Four uniquely regulated pathways were observed in the male high group, namely: metallopeptidases; familyM32; carboxypeptidase Taq family, NtrC family; DctB – DctD, histidine protein kinase; cell cycle family, and NtrC family; GInL – GinG (Fig. 5 B). Two pathways were shown to be uniquely regulated in the female high group, namely: histidine protein kinase; LuxR family, and LuxR family; TtrS – TtrR (Fig. 5 B). 8 predicted pathways were shown to be shared between both the male low and male high treatment group, namely: glycan extension; glycolipid, oxidoreductases; 1.13 acting on oxygenases, metallopeptidases; familyM42; glutamyl amino peptidase family, major facilitator superfamily (MFS); unknown transporters, polyketide synthase (PKS); type-II PKS, polyketide synthase; component type, terpene biosynthesis; squalene/phytoene synthase, and polyketide tailoring proteins; beta-branched incorporating proteins (Fig. 5 B). Two predicted pathways were shown to be significantly regulated in the male low, male high, and female low group, namely: type V secretion system; auto transporter – 1(AT – 1) family, and the major facilitator superfamily (MFS); lipid transporters (Fig. 5 B). Four predicted pathways were differentially regulated in the male low, male high, and female high groups, namely: polysaccharide; structural polysaccharide, CD (cluster of differentiation) molecules, threonine peptidases; familyT3; gamma-glutamyl transferase family, and the metallopeptidases; familyM23 (fig. 5 B). There was one pathway that was significantly regulated in all four treatment groups, the non-ribosomal peptide synthetase (NRPS) and iterative NRPS (fig. 5 B). The relative number of significantly regulated predictive L3 pathways was higher in the male treatment groups compared to the female treatment groups (Fig. 5 B).

### Predictive functional analysis (modules)

Significantly regulated modules were analyzed from the metagenomic shotgun sequencing process based on methods established by Diversigen (New Brighton, MN). In total, 20 significantly regulated modules were found among the four treatment groups. 19 modules were significantly regulated in the male group: 11 modules were downregulated that were associated with methane oxidation, dicarboxylate transport, amino acid transport, nitrogen regulation, betaine biosynthesis, lysine degradation, carotene biosynthesis, microcin C transport, and manganese transport (Fig S3). 8 modules were upregulated in the male group associated with phosphorylated sugar transport, vitamin B12 transport, glutathione biosynthesis, and amino acid transport (Fig S3). 6 modules were significantly regulated in the female group: 4 modules were downregulated that were associated with phosphorylated sugar transport, vitamin B12 transport, and amino acid transport (Fig S3). 2 modules were upregulated that were associated with tetrathionate respiration and phosphorylated mannose transport (Fig. S3).

A Venn diagram was utilized to visualize the unique and shared modules among the four treatment groups (Fig. 5 C). 8 uniquely regulated modules were associated with the male-high group that was associated with dipeptide and dicarboxylate transport, methane oxidation, PTS system, trimethylamine transport, and glycine/betaine/proline transport. There was only one module that was uniquely regulated in the female-high group, namely the tetrathionate respiration two-component regulatory system.

6 significantly regulated modules were shared between the male-low and the male-high groups, namely Microcin C transport, glutathione biosynthesis, lysine degradation, betaine biosynthesis, beta-carotene biosynthesis, and manganese transport. Five differentially regulated modules were shared among the male-low, male-high, and female-high groups, namely the PTS systems associated with cellobiose, mannose, and lactose, as well as the putative polar amino acid transport and Vitamin B12 transport systems (Fig. 5 C). The relative number of significantly regulated modules was higher within the male exposure group compared to the female exposure group (Fig. 5 C).

### Serum untargeted metabolomics

Untargeted metabolomics was also conducted on MPs-treated male and female murine serum. One-way ANOVA in conjunction with Tukey’s post-hoc test with a p-value of 0.05 was utilized. An FDR correction -log(p) value of 0.1 was used to filter additional statistically insignificant data points from the data set. Untargeted metabolomic LC-MS/MS data analysis from murine blood serum showed that 6 significant enzymatic pathways were differentially regulated in the male low group, namely: lysine degradation, amino/nucleotide sugar metabolism, starch/sucrose metabolism, glycine; serine; and threonine metabolism, glyoxylate and dicarboxylate metabolism, and fructose/mannose metabolism (Fig. 6 A). Lysine degradation was the only pathway that was significantly downregulated regulated within the male low group while the rest of the pathways were shown to be upregulated. Only one pathway was shown to be differentially regulated in the male high group, namely: lysine degradation (Fig. 6 B). What was notable was that lysine degradation was significantly downregulated in both male low and male high-exposure groups (fig. 6 A-B).

**Figure 6.**
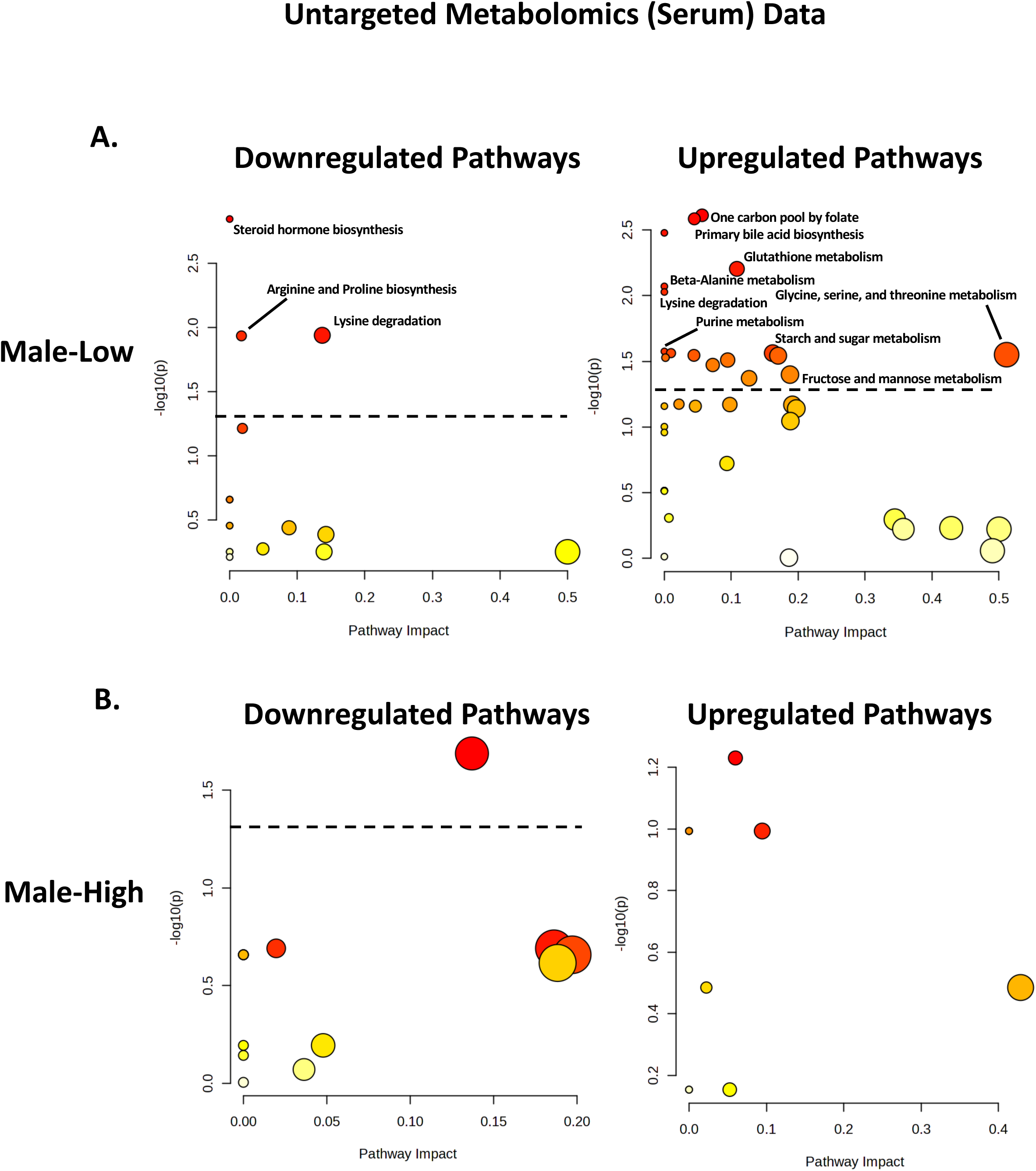
**A.** Untargeted metabolomics conducted on mice blood serum found that low exposure to MPs resulted in a higher number of significantly regulated pathways compared to high MPs exposure. More of these significant pathways were shown to be upregulated during male low MPs exposure. **B.** Untargeted metabolomics conducted on mice blood serum found that high exposure to MPS resulted in a lower number of significantly regulated pathways compared to low MPS exposure in male mice. Only one pathway was significantly downregulated in the male-high group, which was lysine degradation.

**Figure 7.**
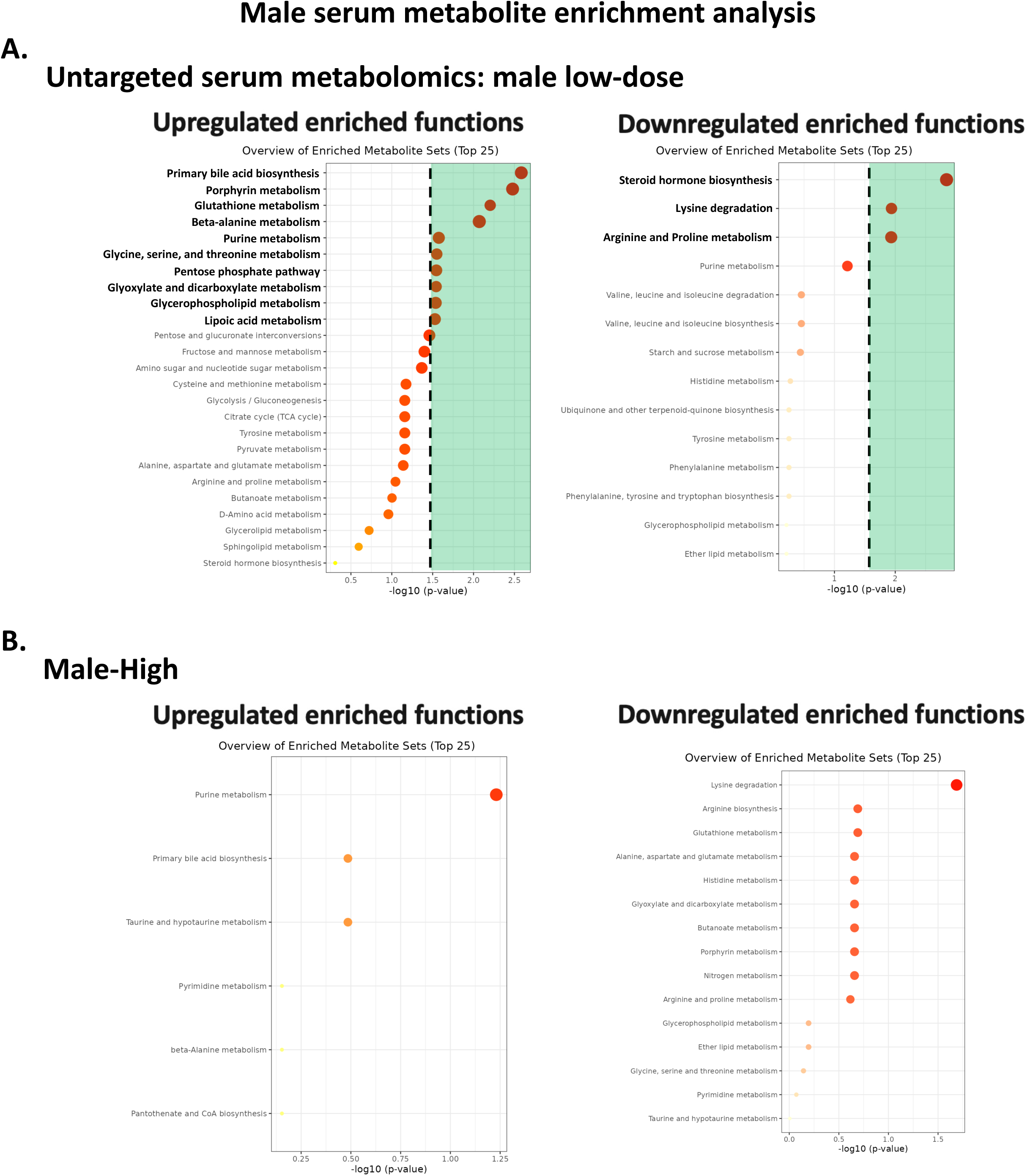
**A.** Pathway enrichment for the low MPs dosage group saw a greater number of upregulated metabolites. **B.** Pathway enrichment for the high MPs dosage group saw no significantly regulated metabolite.

13 significant enzymatic pathways were differentially regulated in the male-low group. 3 pathways were significantly downregulated, namely: steroid hormone biosynthesis, arginine and proline biosynthesis, and lysine degradation (Fig.6 A). 10 pathways were significantly upregulated, namely: primary bile acid biosynthesis, porphyrin metabolism, glutathione metabolism, beta-alanine metabolism, purine metabolism, glycine serine and threonine metabolism, pentose phosphate pathway, glyoxylate and dicarboxylate metabolism, glycerophospholipid metabolism, and lipoic acid metabolism (Fig.6 A). No differentially regulated pathways were observed within the male-high group.

There were significantly more differentially regulated pathways within the female group, most of which were significantly upregulated during both a low and high dose of MPs. 24 differentially regulated pathways were observed in the female-low group. 2 of the pathways were significantly downregulated, namely: d-amino acid metabolism and histidine metabolism (Fig. 8 A). 22 of the pathways were significantly upregulated during low MPs exposure, namely: lysine degradation, valine leucine and isoleucine biosynthesis and degradation, glutathione metabolism, pantothenate and CoA biosynthesis, glyoxylate and dicarboxylate metabolism, pyrimidine metabolism, beta-alanine metabolism, lipoic acid metabolism, glycine serine and threonine metabolism, primary bile acid biosynthesis, d-amino acid metabolism, sphingolipid metabolism, alanine aspartate and glutamate metabolism, cysteine and methionine metabolism, arginine biosynthesis, butanoate metabolism, nitrogen metabolism, terpenoid backbone biosynthesis, and histidine metabolism (Fig. 8 A). 16 pathways were differentially regulated during high MPs exposure, all of which were significantly upregulated, namely: arginine and proline metabolism, arginine biosynthesis, glutathione metabolism, glyoxylate and dicarboxylate metabolism, histidine metabolism, butanoate metabolism, porphyrin metabolism, nitrogen metabolism, alanine aspartate and glutamate metabolism, arachidonic acid metabolism, valine leucine and isoleucine biosynthesis and degradation, glycine serine and threonine metabolism, d-amino acid metabolism, sphingolipid metabolism, and cysteine and methionine metabolism (Fig.8 A).

**Figure 8.**
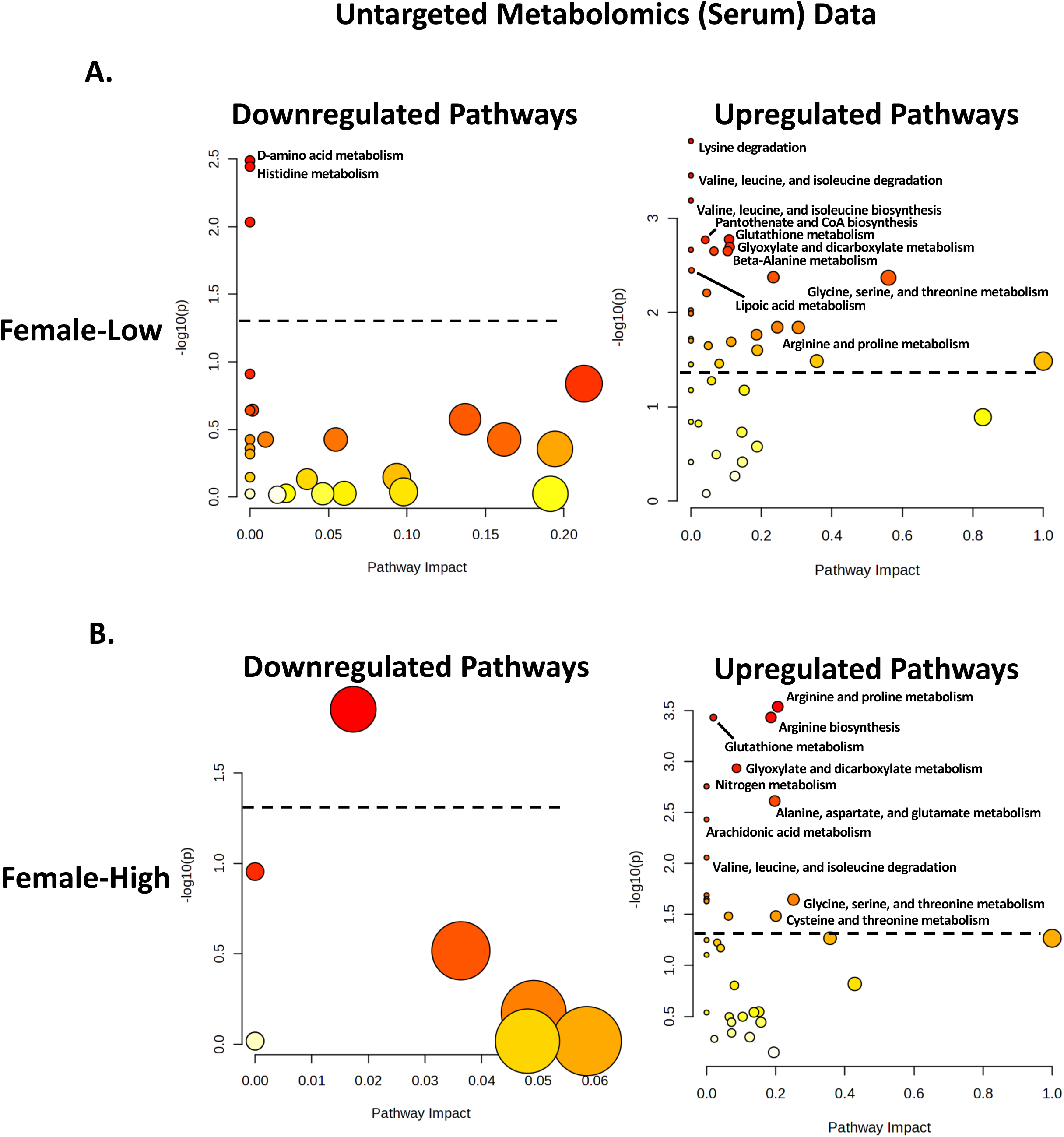
**A.** Dot plots representing the detected pathways associated with the murine blood serum untargeted metabolomics data set, significance was established at p = 0.05 and FDR of x<0.1. We can see that the female-low group had no significant downregulated pathways; however, there are 11 significantly upregulated pathways. **B.** Dot plots representing the detected pathways associated with the murine blood serum untargeted metabolomics data set for the female-high group. Significance was established at p = 0.05 and a pathway impact of x>0.1. We can see that the female-high group had no significant downregulated pathways; however, there are 8 significantly upregulated pathways.

**Figure 9.**
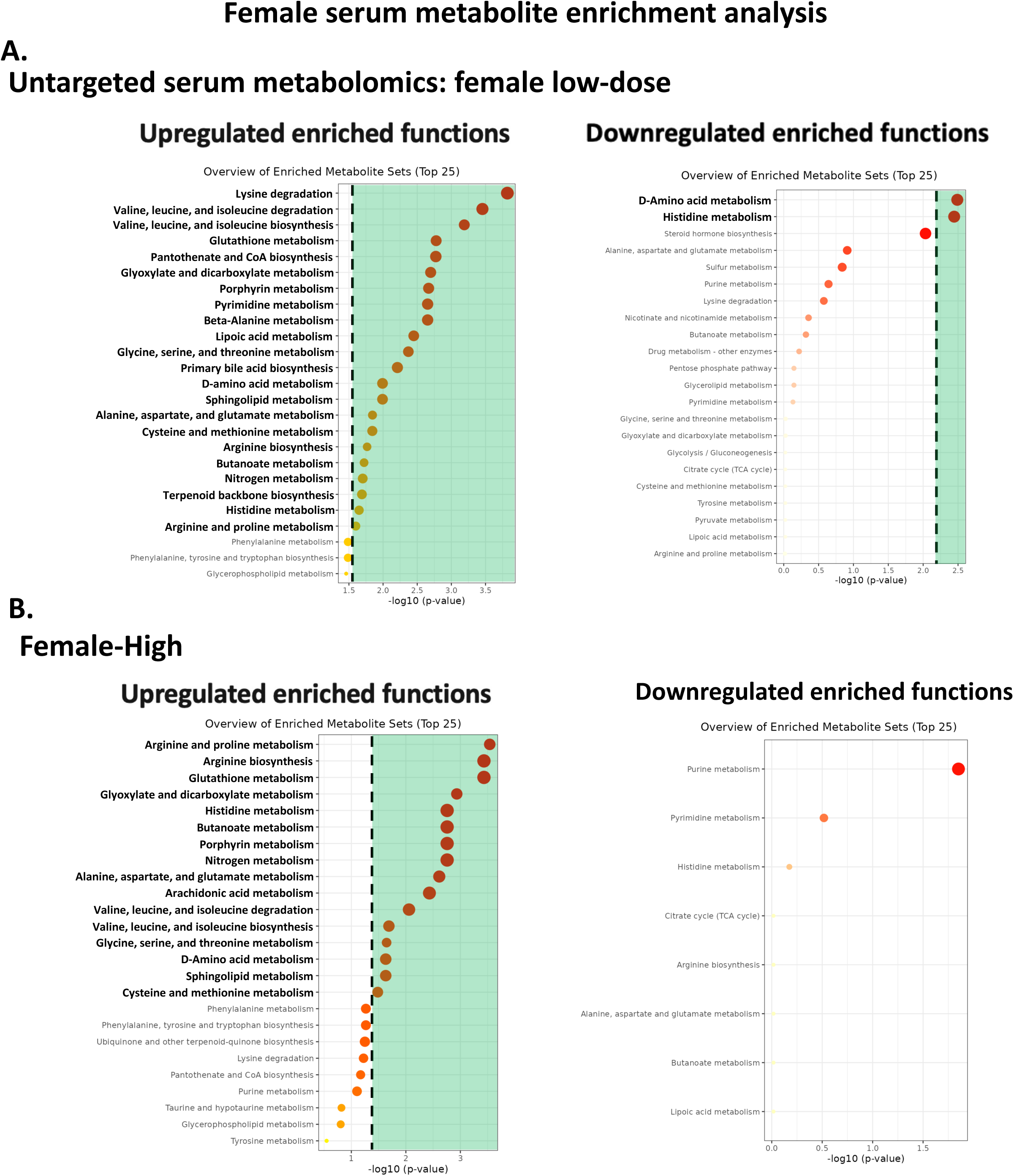
**A.** Pathway enrichment for the female low MPs dosage group saw a greater number of upregulated metabolites. **B.** Pathway enrichment for the high MPs dosage group saw fewer significantly regulated metabolites.

### Serum SCFAs

Specific bacteria known to produce neuroprotective SCFAs were plotted on a relative abundance bar plot from the male mice group. Both the relative abundance of *Akkermansia muciniphila* and *Bacteroides thetaiotaomicron,* bacteria known to produce the neuroprotective SCFAs butyrate, propionate, and acetate, showed a significant decrease in their relative abundances as we increased the dose of MPS exposure in the male mice gut microbiome (Fig. 10 A).

**Figure 10.**
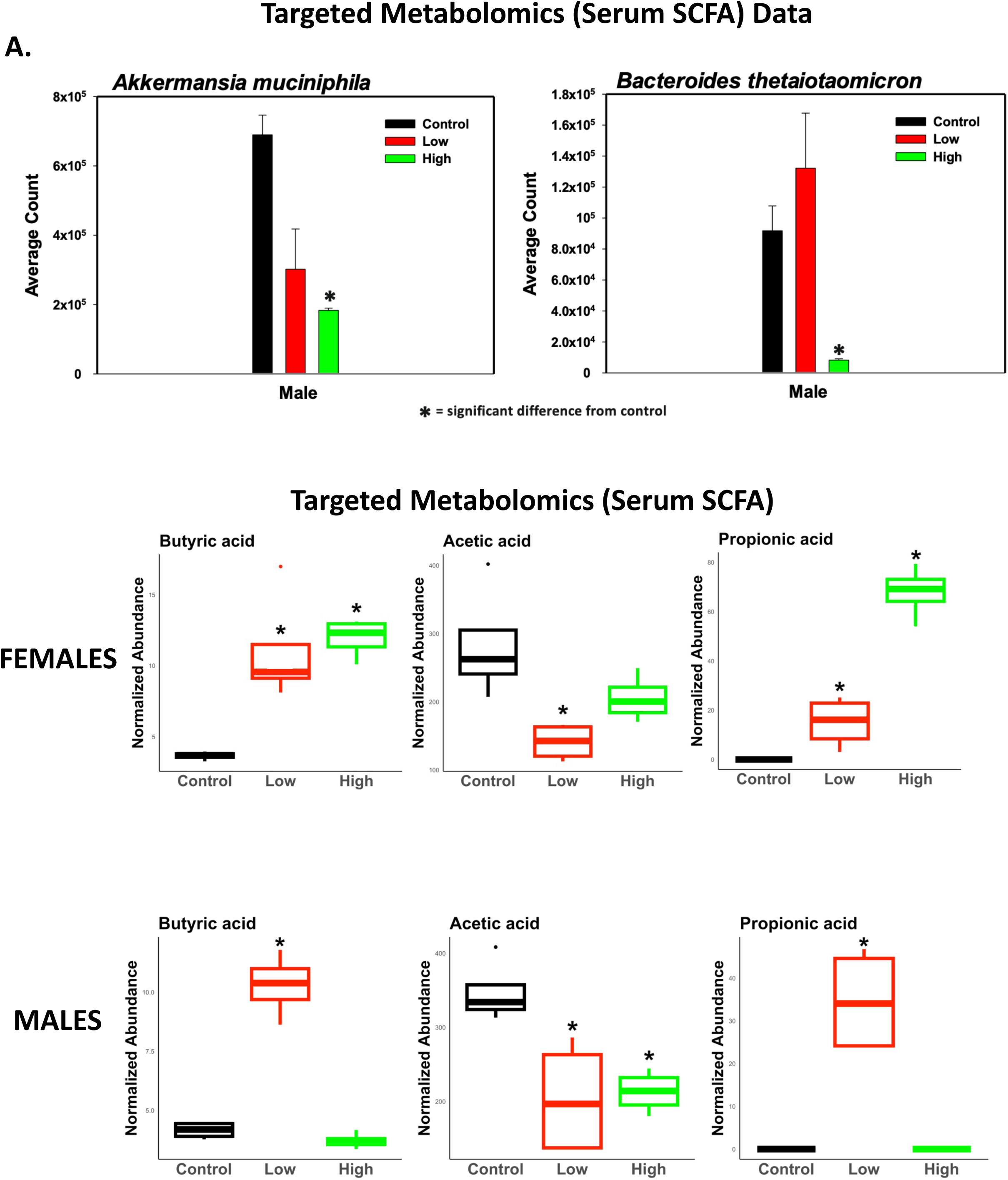
**A.** Averaged raw counts of both Akkermansia muciniphila and Bacteroides thetaiotaomicron (both bacteria known to produce neuroprotective SCFA) were shown to be significantly decreased in male mice at a high MPS dose. **B.** Box plots showing the normalized abundance of three major neuroprotective SCFA: butyrate, acetate, and propionate in the female group. We can see that butyrate and propionate had a significant increase during MPS exposure in a dose-dependent manner while acetate saw a significant decrease at a low MPS exposure. **C.** Box plots showing the normalized abundance of three major neuroprotective SCFA: butyrate, acetate, and propionate in the male group. We can see that butyrate and propionate had a significant increase during MPS exposure while acetate saw a significant decrease at a high MPS exposure.

GC-MS was conducted on murine blood serum samples for both the male and female exposure cohort. Neuroprotective SCFAs and their intermediate metabolites were the focus of our GC-MS measurements. The SCFAs butyrate and propionate were shown to increase in their normalized abundance as we increased the MPS dose, while acetate was shown to be decreased in its normalized abundance within both male and female groups (Fig. 10 B-C).

### Pearson’s Correlation Analysis

Pearson’s correlation analysis was conducted on the normalized SCFA serum concentrations of the male-high group with significantly altered gut-microbial species. Two groups of gut-microbes were established based of anti-inflammatory properties and SCFA production capabilities (Fig. 11 A-B). Both SCFA and microbiome datasets were normalized using the centered log means (CLR) normalization and significance was measured at p-value of 0.05. Benjamani-Hochberg (BH) p-value correction was utilized to control for false discovery rates.

**Figure 11.**
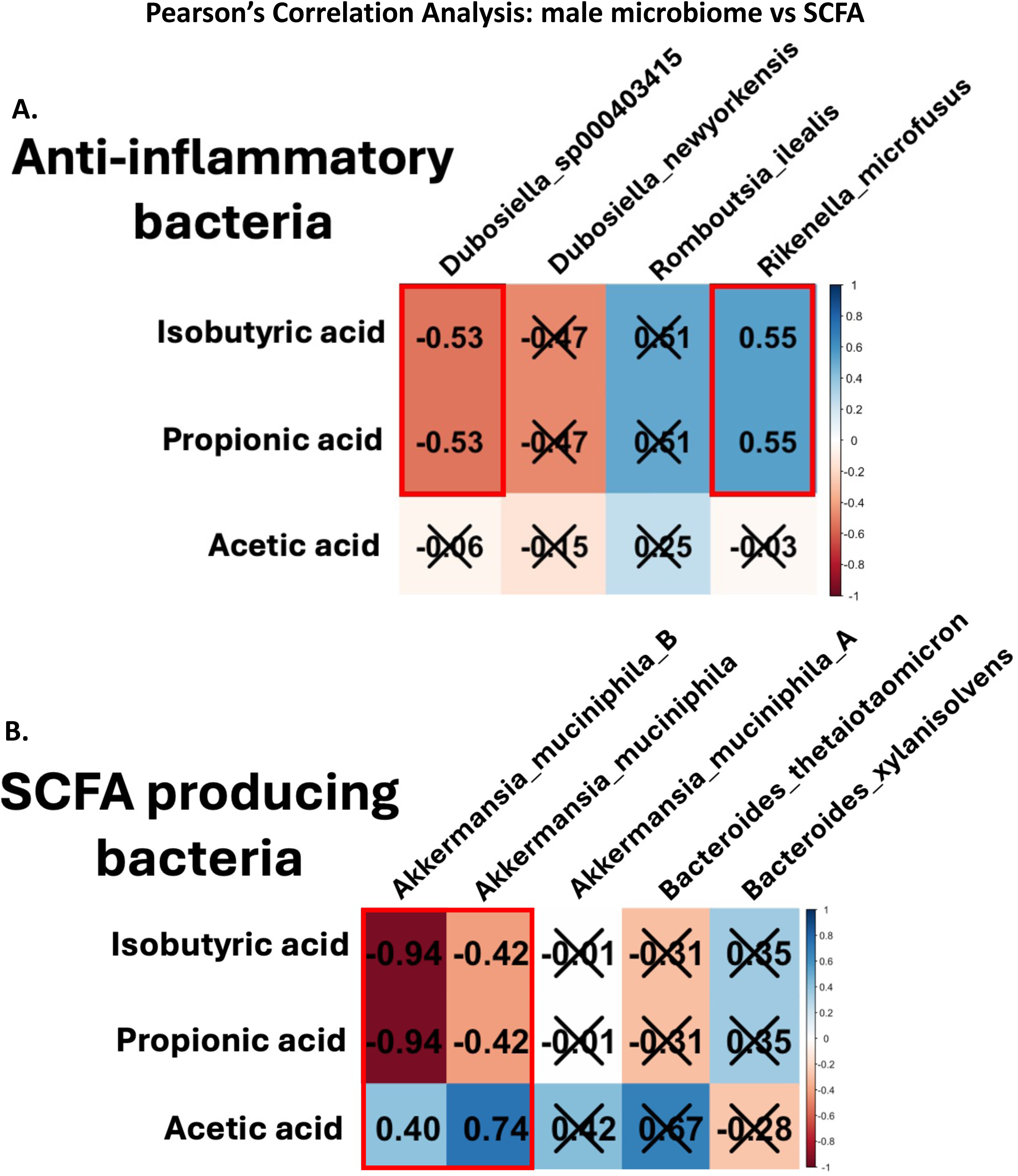
**A.** Pearson’s correlation analysis of normalized serum SCFA concentrations (ug/ml) and relative abundance count of significant anti-inflammatory gut microbes in male-high mice. A significant negative correlation is observed between *Dubosiella_so000403415* with both isobutyric acid and propionic acid serum concentrations. A significant positive correlation was observed with *Rikenella_microfusus* with both isobutryic acid and propionic acid serum concentrations. **B.** Pearson’s correlation analysis of normalized serum SCFA concentrations (ug/ml) and relative abundance count of significant SCFA producing gut microbes in male mice. A significant negative correlation is observed for both *Akkermansia_muciniphila_B* and *Akkermansia_muciniphila* with both isobutyric acid and propionic acid serum concentrations. A positive correlation is observed in both *Akkermansia_muiniphila_B* and *Akkermansia_muciniphila* with acetic acid serum concentrations.

**Figure 12.**
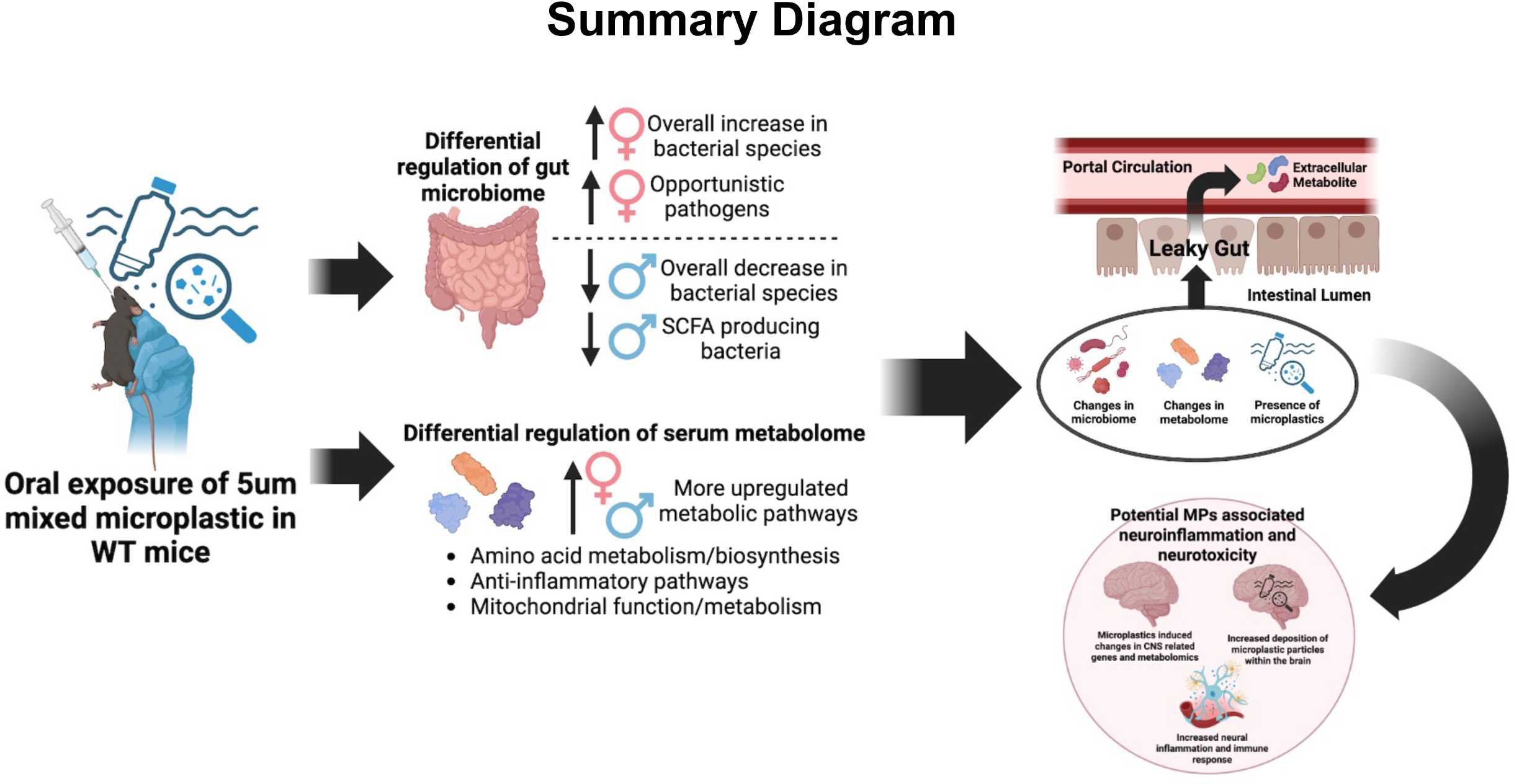
**A.** Metagenomic shotgun sequencing confirmed the presence of a significantly altered gut microbiome at both doses of 5um mixed microplastics within C57BL6 mice. The predictive analysis obtained from the metagenomic shotgun sequencing process showed significant changes to enzymes associated with bacterial carbohydrate metabolism and possible epigenetic markers. Serum untargeted metabolomics confirmed differential changes to enzymes relating to both carbohydrate metabolism and epigenetic pathways. Targeted SCFA metabolomics showed decreases in neuroprotective SCFA in males and increases in females. Overall, the changes associated with the MPs exposure led us to conclude that possible implications may exist within the gut-brain axis of mice in the presence of MPs.

A significant negative correlation was observed between *Dubosiella_so000403415* with both isobutyric acid and propionic acid serum concentrations. A significant positive correlation was observed with *Rikenella_microfusus* with both isobutryic acid and propionic acid serum concentrations (11 A). Pearson’s correlation analysis of normalized serum SCFA concentrations (ug/ml) and relative abundance count of significant SCFA producing gut microbes in male mice. A significant negative correlation is observed for both *Akkermansia_muciniphila_B* and *Akkermansia_muciniphila* with both isobutyric acid and propionic acid serum concentrations. A positive correlation is observed in both *Akkermansia_muiniphila_B* and *Akkermansia_muciniphila* with acetic acid serum concentrations (11 B).

## Discussion

The primary route of human exposure to microplastic particles is oral ingestion (Cox et al. 2019). Microplastics have been detected in various environmental matrices, including food products, soil, and water, underscoring the ubiquitous nature of these contaminants and their interaction with all living organisms (Kosuth et al. 2018; Fan et al. 2023; Oliveri et al. 2020). While extensive research has focused on characterizing the diversity of microplastic congeners in the environment, relatively less is known regarding their impact on the gut microbiome and the associated metabolome (Huerta et al. 2017). Notably, studies have demonstrated that 5 μm polystyrene microspheres can damage the gut epithelium and alter the gut microbiome composition and associated metabolome in male mice (Lu et al 2018). Our study recapitulated the notion that microplastic particles significantly altered the gut microbiome composition and associated metabolome of both male and female mice. However, environmental microplastic exposure is not limited to a single composition or particle size (Alfaro-Nunes et al. 2021). Moreover, the existing studies have predominantly been conducted in male mice, leaving the effects of microplastics on female physiology largely unexplored. To address these important gaps, our research is among the first to utilize a mixed microplastic dose while also utilizing both male and female mice.

Although the overall relative richness of the microbiome (alpha diversity) was not changed among treatment and sex, the distinct group characteristics were significantly different between the high MPs dose and the control group in males. Many factors influence the composition of the gut microbiome, including sex, exposure, diet, and age (Herzog et al. 2021; Shobeiri et al. 2022; Conlon et al. 2014). Here, there were similar numbers of differentially regulated microbial species detected in both sexes. However, we observed a strong overall decrease in bacterial species in the male group, whereas a strong increased pattern was observed within the female treatment groups. Microplastics downregulated probiotic species such as *Akkermansia muciniphila* (Zhang et al. 2021; Ghotaslou et al. 2023) and *Bacteroides thetaiotaomicron* (Delday et al. 2019) in males. Probiotic bacterial species have been shown to have inverse correlations to microplastic-induced pathologies. A study performed in mice reported that probiotic supplementation alleviated the inflammatory effects of polystyrene microparticles and associated reproductive toxicity (Zhang et al. 2023). Such a decrease in probiotic species of bacteria such as *Akkermansia* within the male gut microbiome may be a biomarker of elevated risk of metabolic disorder within the murine gut (Zhang et al. 2021). Microplastics upregulated microbial species within females, some of which were opportunistic pathogens such as *Enterococcus faecalis* (Shah et al. 2024) and *Stenotrophomonas pavanii* (Fluit et al. 2022).

One such explanation for this stark difference can be the fact that male and female mice have different innate biological homeostasis of hormones or other systems that could cause this difference (Valeri et al. 2019). Most of the significantly regulated microbes were common denizens of the murine microbiome; however, a handful of known probiotic species in the genus *Akkermansia, Adlercreutzia, Dubosiella, Bacteroides,* and *Romboutsia* as well as opportunistic pathogens like *Enterococcus faecalis and Stenotrophomonas pavanii* were shown to be significantly regulated during MPS exposure, thus allowing us to say that a significant MPS modulatory effect upon the murine gut microbiome was observed (Fiore et al. 2019; Liu et al. 2023). As mentioned, certain species of bacteria known to produce neuroprotective SCFAs were shown to be downregulated in the male group while opportunistic pathogens saw an increase in relative abundance in both sexes (Fiore et al. 2019; Rodrigues et al. 2022; Wrzosek et al. 2013; Guo et al. 2022).

Untargeted metabolomics data conducted on murine serum samples showed that both male and female groups had significantly altered pathways. More significant pathway hits were seen at a lower dose of MPs in both the male and female groups. This dose-independent pattern may be due to the innate biological systems of both male and female mice compensating for a foreign xenobiotic (MPs) being introduced. Most significant pathways were associated with amino acid, sugar/starch metabolism, and epigenetic markers in both male and female mice. The gut microbiome is known to modulate both starch/sugar and amino acid metabolisms and a homeostasis of these metabolic processes is a biomarker of a healthy gut environment (Chen et al. 2024; Wu et al. 2021). Disruption of amino acid and glucose metabolism has been associated with diseases such as Crohn’s Disease and Irritable Bowel Disease (Scoville et al. 2018; Wu et al. 2021; Capristo et al. 1999) thus, leading us to predict that the presence of sugar and amino acid metabolic imbalance that we observed during MPs exposure may be sign of negative MPs induced biomarker within the gastrointestinal system.

Untargeted metabolomics data from both the liver and pre-frontal cortex from mice of the same cohort showed similar patterns to that of the blood serum untargeted metabolomics. Certain pathways such as beta-alanine metabolism were shown to be significantly regulated in both the liver and serum metabolomics data while certain pathways such as arginine and proline metabolism were seen to be regulated in the serum of the high-dose group and in brain metabolomics of the low-dose (Garcia et al. 2024).

The targeted short-chain fatty acids (SCFAs) metabolomics data set showed that essential fatty acids such as propionate, acetate, and butyrate were significantly regulated within both the male and female groups. SCFAs are produced by the bacterial metabolism of indigestible starch and dietary fibers (Fusco et al. 2023). Certain SCFAs such as butyrate have been shown to regulate tissue inflammation, promote regulatory T-cells, and enhance the production of mucus from intestinal goblet cells (Goldsmith and Sartor 2014; Koh et al. 2020). Acetate has been connected to enhance cardiac health while propionate is a biomarker for a healthy gut microbiome and associated metabolome (Nogal et al. 2021; Reichardt et al. 2014).

Butyric acid and propionic acid were shown to be significantly increased in both male and female murine blood serum; however, acetic acid was shown to be decreased. Two major bacteria species known to produce propionate: *Akkermansia muciniphila* and *Bacteroides thetaiotaomicron* were shown to be decreased in their relative abundance in male mice (Rodrigues et al. 2022; Wrzosek et al. 2013). There are intrinsic host factors that may also play a role in the fluctuation of the SCFA relative abundances. A possible explanation of the difference between the targeted metabolomics and the rest of the data set may be the inherent difference in the biological compartments (feces vs serum). Another possible explanation for these stark patterns is that the excreted (feces) microbiome may differ from the microbiome composition within the intestinal contents.

Due to the inherent nature of the circulatory system being connected to every organ system within a biological entity, it is also reasonable to suspect that there will be systemic effects of MPS exposure in organ compartments throughout the human body. Microplastics could potentially affect the gut-brain axis, influencing communication between the gut and the brain and potentially contributing to neurological disorders such as depression, anxiety, and autism (Xu et al. 2021; Li et al. 2024; Zaheer et al. 2022).

In summary, microplastics altered the murine gut microbiome composition in a sex and dose-dependent manner. For example, microplastics decreased SCFA-producing bacterial species such as *A. muciniphila* and *B. thetaiotaomicron* in males, whreas increased opportunistic pathogens such as *E. faecalis* (Shah et al. 2024) and *S. pavanii* (Fluit et al. 2022) in females. In addition, untargeted metabolomics unveiled MPs-associated dysregulation of the serum metabolic pathways, with more altered by the low MPs dose and female sex status compared to males. Most of the MPs-regulated metabolic pathways were involved in amino acid metabolism and inflammation. Importantly, targeted serum SCFA metabolomics displayed similar patterns in the relative abundances of acetate, butyrate, and propionate in both sexes. Mixed microplastic exposure, at a higher dose, increased the relative level of butyrate and propionate but decreased the level of acetate in both male and female. Taken together, our findings suggest that microplastic exposure can dysregulate the gut microbiome and the associated serum metabolome, contributing to potential negative health impacts relating to metabolic dysregulation, making microplastics an environmental contaminant of public health importance.

## Figure Legends

**Figure S1:** Figure S1. Heatmaps showing the differentially regulated functional enzymes stratified by both sex and treatment. Overall, the male group saw a greater number of differentially regulated functional enzymes than the female group. A dose dependent trend was also seen with regards to the number of differentially regulated functional enzymes detected.

**Figure S2:** Figure S2. Heatmaps showing the differentially regulated L3 functional enzyme pathways stratified by both sex and treatment. Male mice has a greater number of differentially regulated L3 functional enzyme pathways compared to female mice.

**Figure S3:** Figure S3. Heatmaps showing the differentially regulated enzyme modules in MPS exposed mice stratified by both sex and treatment. Male mice (much like that of the enzyme L3 pathways data set) has more differentially regulated enzyme modules compared to female mice.

## Acknowledgments

Funding was supported in part by the National Institutes of Health (NIH) through NIH grant no. R01 ES032037 (E.F.C.), in part by the UW Environmental Pathology/Toxicology Training Program training grant (T32ES007032) (Kyle Joohyung Kim), the Environmental Health and Microbiome Research Center (EHMBRACE), and the Sheldon Murphy Endowment (Cui), UW EDGE Center (NIEHS P30 grant: 5P30ES007033-27); NIH 1U01AG088557, NIH 1R01AG070776

## Author Contributions

KJK is the first author of the manuscript. MG, AR, YJ, and JC are contributing co-authors of the manuscript. EFC, MJC, and HG are contributing authors. JRR and JYC are co-mentors and co-contributors to KJK and the manuscript. EFC and JYC experimental design

