## Supplementary figures and images for "*In vivo* exposure of mixed microplastic particles in mice and its impacts on the murine gut microbiome and metabolome"

### Supplemental figure 1

#### Figure S1

#### Enzymes Data

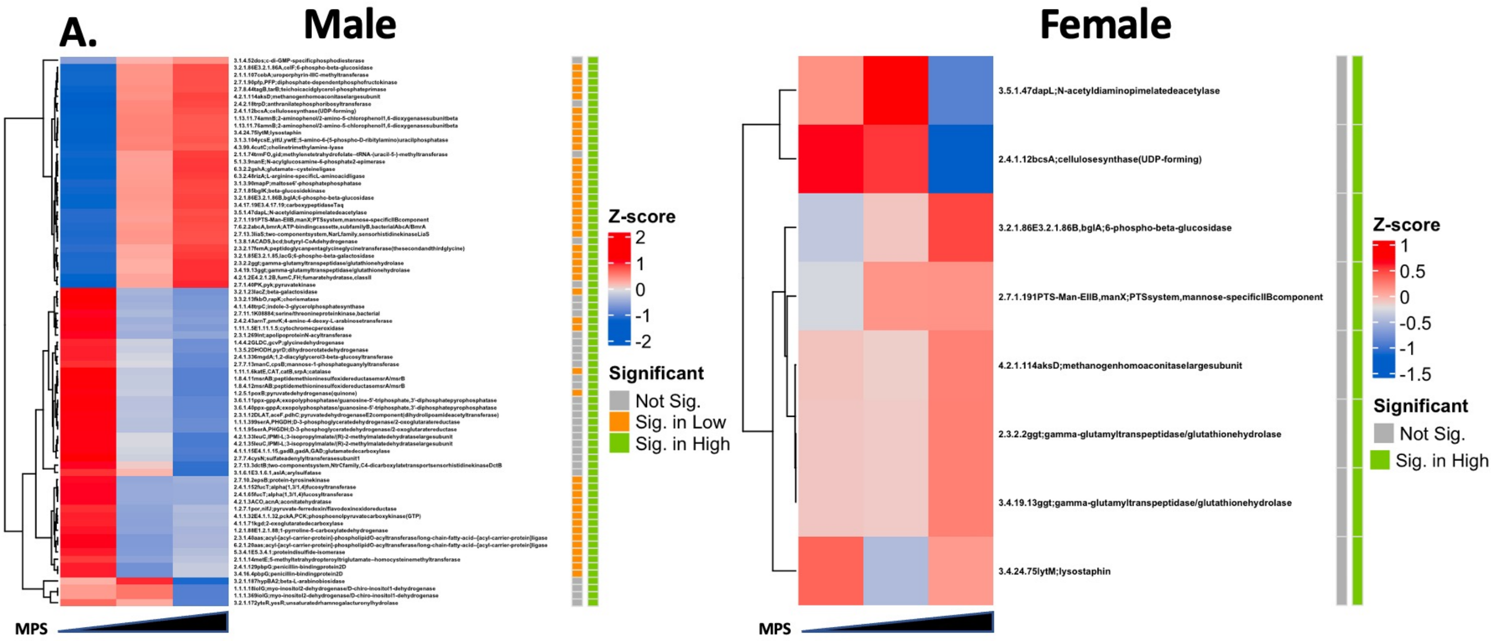

### Supplemental Figure 2

#### Figure S2

##### L3 pathways Data

**B.**

###### Male

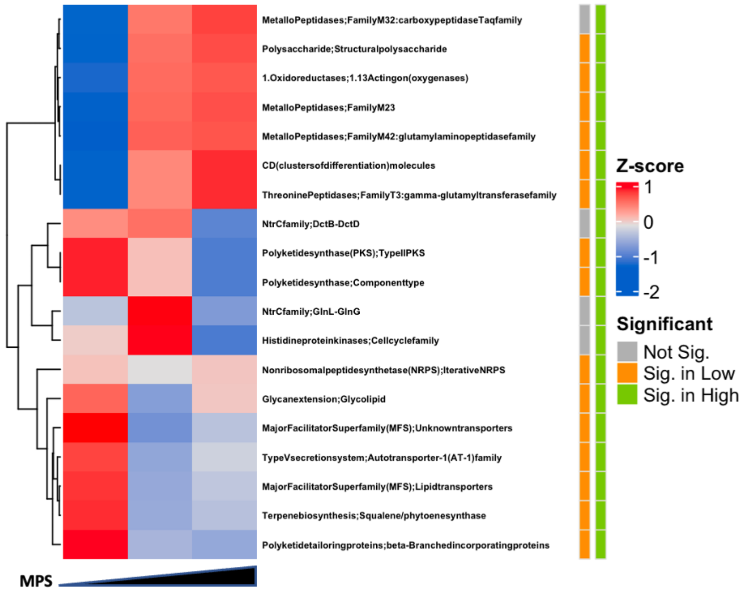


###### Female

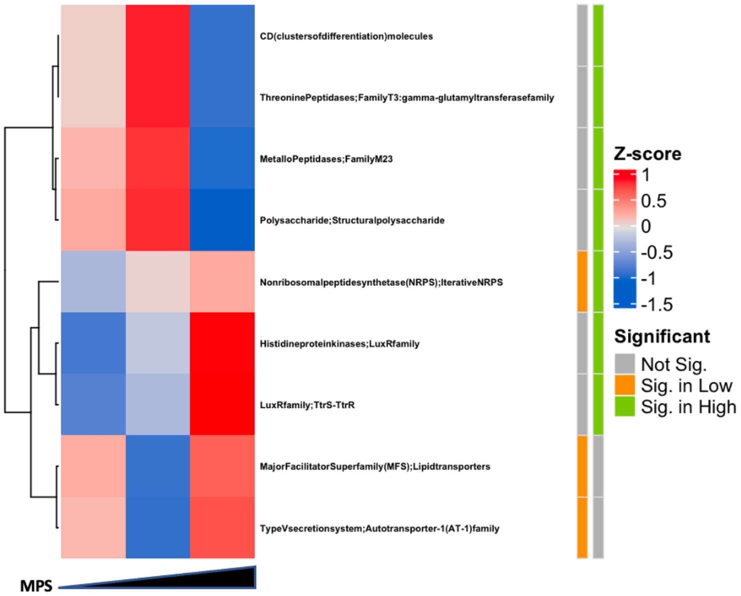

### Supplemental Figure 3

#### Figure S3

## C.

##### Metagenomic predictive pathway modules

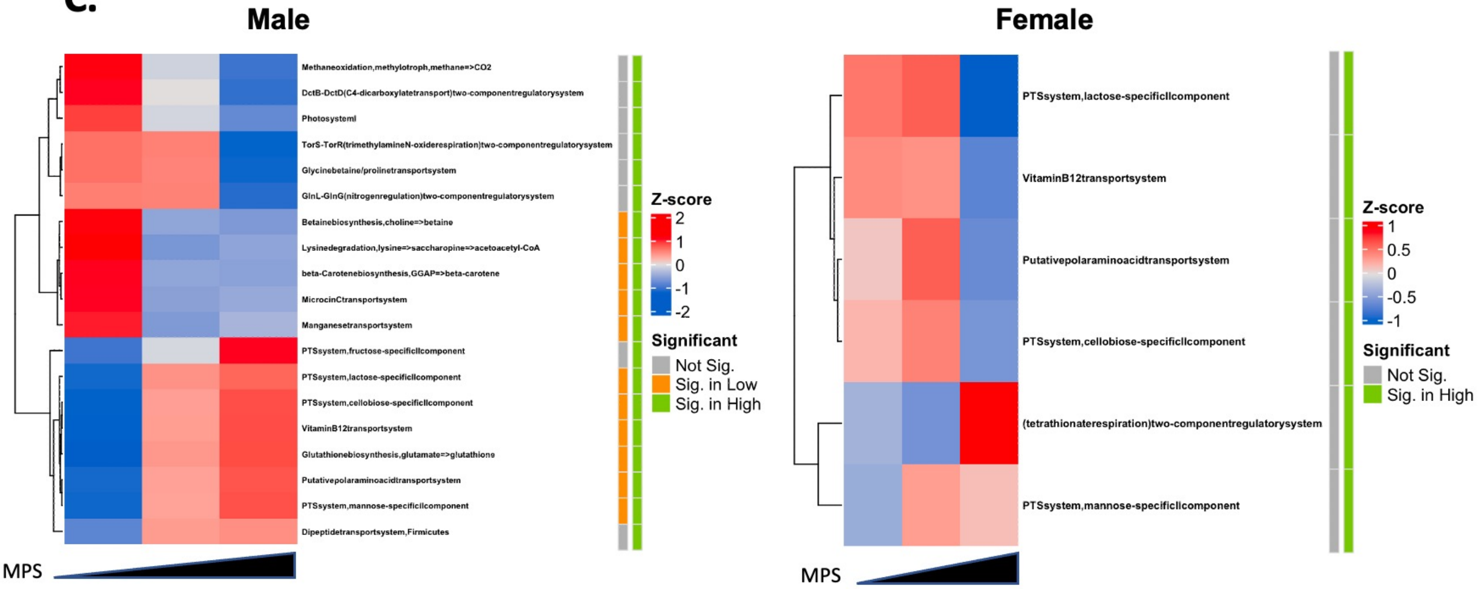
