## Supplemental materials for "*In vivo* exposure of mixed microplastic particles in mice and its impacts on the murine gut microbiome and metabolome"

### Figure S1: Predictive functional analysis (enzymes)

The abundance of the predictive functional microbial enzymes was calculated from the metagenomic shotgun sequencing analysis. MPs exposure in the male group resulted in 76 differentially regulated enzymes: 32 enzymes were significantly upregulated, namely: 3.1.4.52dos;c-di-GMP-specific phosphodiesterase, 3.2.1.86E3.2.1.86A,colF:6-phospho-beta-glucosidase, 2.1.1.107 cobA;uroporphyrin-III C-methyl transferase, 2.7.1.90pfp.PFP:diphosphate-dependent phosphofructokinase, 2.7.8.44tagB.tarB;teichoic acid glycerol-phosphate primase, 4.2.1.114aksD;methanogen homo aconitase large sub unit, 24.2.18trpD;anthranilate phosphoribosyl transferase, 2.4.1.12bcsA;cellulose synthase (UDP-forming), 1.13.11.74amnB;2-amino phenol/ 2-amino-5-chloro phenol 1,6-dioxygenase sub unit beta, 1.13.11.76amnB;2-amino phenol/ 2-amino-5-chloro phenol 1,6-dioxygenase sub unit beta, 3.4.24.75lytM;lysostaphin, 3.1.3.104ycsE,yitU,ywtE:5-amino-6-(5-phospho-D-ribityl amino)uracil phosphatase, 4.3.99.4cutC;choline trimethylamine-lyase, 21.1.74trmFO.gid;methylene tetrahydrofolato-tRNA-(uracil-5-)-methyl transferase, 5.1.3.9nanE;N-acyl glucosamine-6-phosphate2-epimerase, 6.3.2.2gshA:glutamate-cystein ligase, 6.3.2.48rizA;L-arginine-specific L amino acid ligase, 3.1.3.90mapP;maltose 6-phosphate phosphatase, 2.7.1.85bg|K;beta-glucoside kinase, 3.2.1.86E3.21.86B,bglA;6-phospho-beta-glucosidase, 3.4.17.19E3.4.17.19;carboxy peptidase Taq, 3.5.1.47 dapL;N-acetyl diamino pimelate deacetylase, 2.7.1.191PTS-Man-EIIB,manX;PTS system.mannose-specific II B component, 7.6.2.2abcA, bmrA;ATP-binding cassette,sub family B,bacterial Abc A/BmrA, 2.7.13. 3liaS:two-component system.Nar L family sensor histidine kinase

Liase, 1.3.8.1ACADS,bcd; butyryl-CoA dehydrogenase, 2.3.2.17lemA;peptidoglycan pentaglycine glycine transferase(the second and third glycine), 3.2.1.85E3.21.85,lacG;6 phospho-beta-galactosidase, 2.3.2.299t:gamma-glutamyl transpeptidase/ glutathione hydrolase, 3.4.19.13ggt:gamma-glutamyl transpeptidase/ glutathione hydrolase, 4.2.1.2E4 2.1.2B,fumC.FH fumarate hydratase, class II, and 2.7.1.40PK.pyki,pyruvate kinase (Fig. 5 A). 44 enzymes were significantly downregulated in the male group, namely: 3.2.1.23lacz;beta-galactosidase, 3.3.2.138kbO,rapK;chorismatase, 41.1.48trpC Indole-3-glycerol phosphate synthase, 2.7.11.1K08884;serine/ threonine protein kinase, bacterial, 24.2.43arnT,pmrK;4-amino -4-deoxy-L-arabinose transferase, 1.11.1.5E1.11.1.5;cytochrome c peroxidase, 2.3.1.269int;apolipoprotein N-acyl transferase, 1.4.4.2GLDC.gcvP:glycine dehydrogenase, 1.3.5.2DHODH.pyrD;dihydro orotate dehydrogenase, 3.4.1.336mgdA; 1,2-diacyl glycerol 3-beta-glucosyl transferase, 2.7.7.13manC,opsB;mannose-1-phosphate guanylyl transferase, 1.11.1.6katE,CAT,catB,srpA;catalase, 1.8.4.11 msrAB;peptide methionine sulfoxide reductase msrA/msrB, 1.8.4.12msrAB;peptide methionine sulfoxide reductase msrA/msrB, 1.2.5.1 poxB;pyruvate dehydrogenase (quinone), 3.6.1.11 ppx-9ppA;exopolyphosphatase/ guanosine-5'-triphosphate, 3*-diphosphate pyrophosphatase, 3.6.1.40ppx-gppA;exopolyphosphatase/ guanosine-5'-triphosphate, 3"-diphosphate pyrophosphatase, 2.3.1.12DLAT,aceF,pdhC;pyruvate dehydrogenase E2 component (dihydro lipoamide acetyl transferase), 1.1.1.399serA,PHGDH;D-3-phosphoglycerate dehydrogenase/ 2-oxoglutarate reductase, 1.1.1.95serA,PHGDH;D-3-phosphoglycerate dehydrogenase/ 2-oxoglutarate reductase, 4.2.1.33leuCIPMI-L;3-isopropyl malate/ (R)-2-methyl malate dehydratase large sub unit, 4.2.1.35leuC,JPMI-L;3-isopropyl malate/ (R)-2-methyl malate dehydratase large sub unit, 4.1.1.15E4.1.1.15,gadB.gadA,GAD:glutamate decarboxylase, 2.7.7 AcysN:sulfate adenylyl transferase sub unit 1, 2.7.13.3dctB;two-component system.NtrCfamily,C4-dicarboxylate transport sensor histidine kinase Dct B, 3.1.6.13.1.6.1,aslA;aryl sulfatase, 2.7.10.2epsB:protein-tyrosine kinase, 24.1.152fucT;alpha (1,3/1,4) fucosyl transferase, 24.1.65fucT; alpha (1,3/1,4) fucosyl transferase,

4.2.1.3ACO,acnA;aconitate hydratase, 1.2.7.1por nifJ;pyruvate-ferredoxin/ flavodoxin oxidoreductase, 4.1.1.324.1.1.32pckA,PCK:phosphoenol pyruvatocarboxy kinase(GTP), 4.1.1.71kgd: 2-oxoglutarate decarboxylase, 1.2.1.881.2.1.88:1-pyrroline-5-carboxylate dehydrogenase, 2.3.1.40 aas :acyl-(acyl-carrier protein]-phospholipid O-acyl transferase/ long-chain fatty acid-(acyl-carrier-protein) ligase, 6.2.1.20eas:acy/-[acyl-carrier-protein/-phospholipid O-acyl transferase/ long-chain-fatty-acid-(acyl-carrier-protein) ligase, 5.3.4.15.3.4.1:protein disulfide-isomerase, 2.1.1.14 metE; 5-methyl tetrahydropteroyl triglutamate – homocysteine methyl transferase, 24.1.129pbpG:penicillin-binding protein 20, 3.4.16.4pbp:penicillin-binding protein 20, 3. 2.1.187hypBA2;beta L-arabinobiosidase, 1.1.1.18iolG:myo-inositol 2-dehydrogenase/ D-chiro-inositol 1-dehydrogenase, 1.1.1.369ioiG;myo-inositol 2-dehydrogenase/ D-chiro-inositol 2-dehydrogenase, and 321.172yteR_ye=R: unsaturated rhamnogalacturonyl hydrolase (Fig. 5 A). 8 differentially regulated enzymes were detected only in the female-high exposure group, of which 2 were significantly down regulated: 3.5.1.47dapL; N-acetyl diamino pimelate deacetylase and 2.4.1.12bcsA; cellulose synthase (UDP forming) (Fig. 5 A). 6 enzymes were significantly downregulated in the female-high group, namely: 3.2.1.86E3.2.1.86B,bglA;6-phospho-beta-glucosidase, 2.7.1.19PTS-Man-EIIB,manX;PTS system,mannose-specific II B component, 4.2.114aksD;methanogen homo aconitase large sub unit, 2.3.2.2ggt;gamma-glutamyl transpeptidase/ glutathione hydrolase, 3.4.19.13ggt;gamma-glutamyl transpeptidase/ glutathione hydrolase, and 3.4.24.75lytM;lysostaphin (Fig. 5 A).

A Venn diagram was utilized to stratify the enzyme data set by both sex and treatment. There were 30 uniquely regulated enzymes within the male-high group, namely 3.1.6.13.1.6.1aslA;aryl sulfatase, 3.2.1.172yteR,yesR;unsaturated rhamnogalacturonyl hydrolase, 2.7.11.1K08884;serine/threonine protein kinase bacterial, 1.3.8.1ACADS,bcd; butyry|-CoA dehydrogenase, 1.4.4.2GLDC.gcvP;glycine dehydrogenase, 2.7.1.40PK,pyk;pyruvate kinase, 1.1.1.18iolG;myo-inositol 2-dehydrogenase/D-chiro-inositol1-dehydrogenase, 1.1.1.369ioG;myo-inositol2-dehydrogenase/D-chiro-inositol1-dehydrogenase, 4.2.1.33LeuC,IPMI-L; 3-isopropyl malate/ (R)-2-methyl malate dehydratase large subunit, 4.2.1.35leuC,IPMI-L;3-isopropylmalate/(R)-2-methylmalatedehydrataselargesubunit, 4.1.1.154.1.1.15,gadB,gadA, GAD;glutamate decarboxylase, 2.1.1.74trmFO,gid; methylene tetrahydro folate-tRNA-(uracil-5-)-methyl transferase, 3.6.1.11ppx-gppA;exopolyphosphatase/ guanosine-5'-triphosphate, 3'-diphosphate pyrophosphatase, 3.6.1.40ppx-gppA;exopoly phosphatase/ guanosine-5'-triphosphate, 3'-diphosphate pyrophosphatase, 2.4.1.336mgdA;1,2-diacylglycerol 3-beta-glucosyl tratisferase, 1.1.1.399serA,PHGDH;D-3-phosphoglycerate dehydrogenase/ 2-oxoglutarate reductase, 1.1.1.95 serA, PHGDH;D-3-phosphoglycerate dehydrogenase/ 2-oxoglutarate reductase, 2.3.1.269lnt;apolipoprotein N-acyl transferase, 2.4.2.18trpD;anthranilate phosphoribosyl transferase, 2.7.7.13manC, cpsB;mannose-1-phosphate guanylyl transferase, 2.7.7.4cysN;sulfate adenylyl transferase subunit 1, 4.1.1.48trpC;indole-3-glycerol phosphate synthase, 2.3.1.12DLAT,aceF,pdhC;pyruvate dehydrogenase E2 component (dihydro lipoamide acetyl transferase), 2.7.13.3dctB;two-component system, NtrCfamily, C4-dicarboxylate transport sensor histidine kinase DctB, 3.1.4.52dos;c-di-GMP-specific phosphodiesterase, 3.3.2.13fkbO,rapK;chorismatase, 1.8.4.11 msrAB;peptide methionine sulfoxide reductase msrA/msrB, 1.8.4.12 msrAB;peptide methionine sulfoxide reductase msrA/ msrB., 1.3.5.2DHODH,pyrD;dihydro orotate dehydrogenase, 3.2.1.187hypBA2;beta-L-arabino biosidase. (Fig 5B.)

38 enzymes were significantly regulated by MPS exposure in both the male-low and the male-high group, namely: 3.2.1.863.2.1.86B, bglA;6-phospho-beta-glucosidase, 4.3.99.4cutC;choline trimethylamine-lyase, 3.1.3.104ycE, vitU, ywtE;5-amino-6-(5-phospho-D-ribityl amino) uracil phosphatase, 7.6.2.2abcA,bmrA;ATP-binding cassette, subfamily, bacterial AbcA/BmrA, 3.2.1.23lacz;beta-galactosidase, 2.4.1.129pbpG;penicillin-binding protein 2D, 3.4.16.4pbpG ;penicillin-binding protein 2D, 1.2.7.1por,nif;pyruvate-ferredoxin/ flavodoxin oxidoreductase, 4.2.1.3ACO,acnA;aconitate hydratase, 1.2.1.88E1.2.1.88;1-pyrroline-5-carboxylate dehydrogenase, 4.2.1.114aksD;methanogen homo aconitase large subunit,

2.7.1.191PTS-Man-EIIB, manX; PTS system, mannose-specific IiB component, 1.11.1.51.11.1.5;cytochrome c peroxidase, 1.13.11.74amnB;2-amino phenol/ 2-amino-5-chloro phenol1, 6-dioxygenase Sub unit beta, 1.13.11.76amnB;2-amino phenol/ 2-amino-5-chloro phenol1,6-dioxygenase sub unit beta, 2.4.1.12bcsA;cellulose synthase (UDP-forming), 3.5.1.47 dapL;N-acetyl diamino pimelate deacetylase, 2.1.1.14metE;5-methyl tetrahydropteroy triglutamate-nomo cysteine methyl transferase, 5.3.4.15.3.4.1;protein disulfide-isomerase, 3.4.17.193.4.17.19;carboxy peptidase Taq, 5.1.3.9nanE;N-acyl glucosamine-6-phosphate 2-epimerase, 6.3.2.48rizA;L-arginine-specific-amino acid ligase, 1.11.1.6katE, CAT, catB, spA; catalase, 2.3.2.2ggt;gamma-glutamyl transpeptidase/ glutathione hydrolase, 3.4.19.13ggt:gamma-glutamyl transpeptidase/ glutathione hydrolase, 3.4.24.75lytM;lysostaphin, 3.2.1.86E3.2.1.86A,celF;6-phospho-beta-glucosidase, 4.1.1.71kgd;2-oxoglutarate decarboxylase, 2.7.10.2epsB;protein-tyrosine kinase, 2.4.2.43arnT,pmrK;4-amino-4-deoxy-L-arabinose transferase, 2.7.8.44tagB ,tarB;teichoic acid glycerol-phosphate primase, 2.7.1.85bg|K;beta-glucoside kinase, 2.1.1.107 cobA;uroporphyrin-III C-methyl transferase, 2.7.13.3liaS;two-component system, Nar L family,sensor histidine kinase lias, 4.1.1.32E4.1.1.32,pckA,PCK;phosphoenol pyruvate carboxy kinase(GTP), 2.3.2.17femA;peptidoglycan pentaglycine glycine transferase(the second and third glycine) (Fig 5B.)

8 enzymes were significantly regulated by MPS exposure in three groups: male-low, male-high, and female-high: 3.2.1.86E3.2.1.86B,bglA;6-phospho-beta-glucosidase, 4.2.1.114aksD; methanogen homo aconitase large sub unit, 2.7.1.191PTS-Man-EIIB,manX; PTSsystem,mannose-specific IiB component, 2.4.1.12bcsA; cellulose synthase (UDP-forming), 2.3.2.2ggt;gamma-glutamyl transpeptidase/ glutathione hydrolase, 3.4.19.13ggt;gamma-glutamyl transpeptidase/ glutathione hydrolase, 3.5.1.47 dapL; N-acetyl diamino pimelate deacetylase, 3.4.24.75lytM; lysostaphin (Fig. 5B).

In both males and females, a dose-response relationship can be seen in that low MPs exposure had a weaker effect on the functional predictive enzyme hits compared to high MPs exposure. Male mice had more significantly regulated enzymes associated with MPs exposure, signifying that the male group was more sensitive toward MPS exposure compared to the female group (Fig. 5C).

**Figure S2: Predictive functional analysis (L3 pathways)**

Significantly regulated L3 predictive pathways were analyzed from the metagenomic shotgun sequencing. In total, 22 significantly regulated L3 predictive pathways were found among the four treatment groups. 19 L3 predictive pathways were significantly regulated in the male group: 10 L3 predictive pathways were downregulated, namely the NtrC family; DctB; DctD, Polyketide synthase(PKS); Type II PKS, Polyketide synthase; Component type, NtrC family; GlnL-GlnG, Histidine protein kinases; Cell cycle family, Major Facilitator Superfamily (MFS); Unknown transporters, Type V Secretion System; Autotransporter-1(AT-1) family, Major Facilitator Superfamily (MFS); Lipid transporters, Terpene biosynthesis; Squalene/phytoene synthase, and Polyketide tailoring proteins; beta-branched incorporating proteins (Fig. 6A). 9 pathways were upregulated during MPs exposure in males, namely Metallopeptidase; familyM32:carboxypeptidase Taq family, Polysaccharide; structural polysaccharide, Oxidoreductase; 1.13 Acting on oxygenase, Metallopeptidases; familyM23, Metallopeptidases; familyM42: glutamyl aminopeptidase family, CD (Clusters of Differentiation) molecules, Threonine Peptidases; familyT3: gamma-glutamyl transferase family, Non ribosomal peptide synthetase (NRPS); iterative NRPS, and Glycan extension; glycolipid (Fig. 6 A). Four pathways were downregulated during MPs exposure in females: CD (Clusters of Differentiation) molecules, Threonine Peptidases; familyT3: gamma-glutamyl transferase family, Metallopeptidases; familyM23, and Polysaccharide: structural polysaccharide while five pathways were upregulated in the female group: Non-ribosomal peptide synthase (NRPS); Iterative NRPS, Histidine protein kinases; LuxR family, LuxR family; Ttrs-TtrR, Major Facilitator Superfamily (MFS); Lipid transporters, and Type V secretion system; Autotransporter-1 (AT-1) family (Fig. 6 A).

A Venn diagram was utilized to observe the unique and shared differentially regulated L3 predictive pathways among the treatment groups. Four uniquely regulated pathways were observed in the male high group, namely: metallopeptidases; familyM32; carboxypeptidase Taq family, NtrC family; DctB – DctD, histidine protein kinase; cell cycle family, and NtrC family; GlnL – GinG (Fig. 6 B). Two pathways were shown to be uniquely regulated in the female high group, namely: histidine protein kinase; LuxR family, and LuxR family; TtrS – TtrR (Fig. 6 B). 8 pathways were shown to be shared between both the male low and male high treatment group, namely: glycan extension; glycolipid, oxidoreductases; 1.13 acting on oxygenases, metallopeptidases; familyM42; glutamyl amino peptidase family, major facilitator superfamily (MFS); unknown transporters, polyketide synthase (PKS); type-II PKS, polyketide synthase; component type, terpene biosynthesis; squalene/phytoene synthase, and polyketide tailoring proteins; beta-branched incorporating proteins (Fig. 6 B). Two pathways were shown to be significantly regulated in the male low, male high, and female low group, namely: type V secretion system; auto transporter – 1(AT – 1) family, and the major facilitator superfamily (MFS); lipid transporters (Fig. 6 B). Four pathways were differentially regulated in the male low, male high, and female high groups, namely: polysaccharide; structural polysaccharide, CD (cluster of differentiation) molecules, threonine peptidases; familyT3; gamma-glutamyl transferase family, and the metallopeptidases; familyM23 (fig. 6 B). Interestingly, there was one pathway that was significantly regulated in all four treatment groups, namely: the non-ribosomal peptide synthetase (NRPS); and iterative NRPS (fig. 6 B). The relative number of significantly regulated predictive L3 pathways was higher in the male treatment groups compared to the female treatment groups (Fig. 6 C).

**Figure S3: Predictive functional analysis (modules)**

Significantly regulated modules were analyzed from the metagenomic shotgun sequencing process. In total, 20 significantly regulated modules were found among the four treatment groups. 19 modules were significantly regulated in the male group: 11 modules were downregulated, namely Methane oxidation; methylotroph; methane=>cO2; DetB-DctD (C4-dicarboxylate transport) two-component regulatory system, Photo system 1, TorS-TorR (trimethylamine N-oxide respiration) two-component regulatory system, Glycine betaine/ proline transport system, GlnL-GlnG (nitrogen regulation) two-component regulatory system, Betaine biosynthesis,choline=>betaine, Lysine degradation, lysine => saccharopine: aceto acetyl-CoA, beta-Carotene biosynthesis, GGAPE>beta-carotene, Microcin C transport system, Manganese transport system (Fig 7. A). 8 modules were upregulated in the male group, namely: PTS system, fructose-specific II component, PTS system, lactose-specific II component, PTS system,cellobiose-specific II component, Vitamin B 12 transport system, Glutathione biosynthesis glutamate=>glutathione, Putative polar amino acid transport system, PTS system mannose-specific II component, and Dipeptide transport system,Firmicute (Fig 7. A). 6 modules were significantly regulated in the female group: 4 modules were downregulated, namely: PTS system,lactose-specific II component, Vitamin B12 transport system, Putative polar amino acid transport system, and the PTS system,cellobiose-specific II system (Fig 7. A). 2 modules were upregulated, namely: (tetrathionate respiration) two-component regulatory system and the PTS system,mannose-specific II component (Fig. 7 A).

A Venn diagram was utilized to observe the unique and shared modules among the four treatment groups (Fig. 7B). 8 uniquely regulated modules were associated with the male-high group, namely M00566 Dipeptide transport system - Firmicutes, M00174 Methane oxidation methylotroph methane=>CO2, M00504 DctB-DctD (C4-dicarboxylate transport) two-component regulatory system, M00304 PTSsystem fructose-specific II component, M00497 GlnL-GlnG(nitrogen regulation) two-component regulatory system, M00163 Photosystem-I, M00455 TorS-TorR(trimethylamine-oxide respiration) two-component regulatory system, and M00208

Glycine betaine/proline transport system. There was only one module that was uniquely regulated in the female-high group, namely the M00514 TtrS-TtrR(tetrathionate respiration) two-component regulatory system.

6 significantly regulated modules were shared between the male-low and the male-high groups, namely M00349 Microcin C transport system, M00118 Glutathione biosynthesis glutamate=>glutathione, M00032 Lysine degradation lysine=>saccharopine=>acetoacetyl-CoA, M00555 Betaine biosynthesis choline=>betaine, M00097 beta-Carotene biosynthesis GGAP=>beta-carotene, and the M00316 Manganese transport system. Five differentially regulated enzymes were shared among the male-low, male-high, and the female-high group, namely the M00275 PTS system cellobiose-specific II component, M00276 PTS system mannose-specific II component, M00281 PTS system lactose-specific II component, M00236 Putative polar amino acid transport system, and the M00241 VitaminB12 transport system (Fig. 7B). The relative number of significantly regulated modules was higher within the male exposure group compared to the female exposure group (Fig. 7C).

Untargeted metabolomic LC-MS/MS data analysis from murine blood serum showed that 6 significant enzymatic pathways were differentially regulated in the male low group, namely: lysine degradation, amino/nucleotide sugar metabolism, starch/sucrose metabolism, glycine; serine; and threonine metabolism, glyoxylate and dicarboxylate metabolism, and fructose/mannose metabolism (Fig. 8 A). Lysine degradation was the only pathway that was significantly downregulated regulated within the male low group while the rest of the pathways were shown to be upregulated. Only one pathway was shown to be differentially regulated in the male high group, namely: lysine degradation (Fig. 8 B). An interesting point is that lysine degradation was significantly downregulated in both male low and male high-exposure groups (fig. 8 A-B).
